# Au_18_(L-NIBC)_14_ targets ASC oligomerization for autoimmune disease treatment

**DOI:** 10.1101/2025.03.25.645153

**Authors:** Kai Yang, Xinyue Zhao, Zhupeng Xiao, Cheng Zeng, Lei Shen, Taolei Sun

## Abstract

Gold nanoclusters (AuNCs), unlike conventional nanoparticles, possess molecular characteristics besides ultrasmall nano-features. Recently, we and others showed that AuNCs are promising in treatments of various major diseases. However, the AuNCs used were usually mixtures, and the specific target and their relationship with AuNC structures are unclear, which largely restrict their druggability. Multiple sclerosis (MS) is an autoimmune disease (AID) implicating central nervous system (CNS), in which drug discovery is challenging. Here we used Au_18_(L-*N*-isobutyryl-L-cysteine)_14_ (Au_18_(L-NIBC)_14_) and Au_25_(L-NIBC)_18_, two AuNCs with the same ligand, to report their much different therapeutic effects in experimental autoimmune encephalomyelitis (EAE). We show that, Au_18_(L-NIBC)_14_, but not Au_25_(L-NIBC)_18_, specifically targets apoptosis-associated speck-like protein containing a CARD (ASC) oligomerization, thus inhibit the activation of ASC-dependent inflammasomes, resulting in comprehensive restoration of cytokine homeostasis in the CNS of EAE mice. Au_18_(L-NIBC)_14_ significantly prevents axon demyelination, protects blood-brain barrier, blocks immune cell infiltration into CNS, and completely prevents motor deficits and relieve the early-cognitive impairments of EAE mice. Remarkable efficacies were also observed in animal models of inflammatory bowel disease, psoriasis, systemic lupus erythematosus, indicating a broad prospect in AIDs treatments. Especially, definite molecular structure, specific target, clear mechanism, and exact therapeutic effects imply a good druggability of Au_18_(L-NIBC)_14_.

## Text

Gold nanoclusters (AuNCs) are one of the forefronts of materials science^1^, which have been extensively studied in catalysis,^2^ drug delivery^3^, bioimaging^4^, photothermal therapy^5^, *etc*. They are ultrasmall nanoparticles with sizes smaller than 3 nm, and, unlike gold nanoparticles larger than 3 nm, possess simultaneously molecular structures^2^ that can be described by mass spectrometer (MS). Recently, we and others showed that AuNCs possess excellent pharmacological activities in various disorders, such as Alzheimer’s disease^6–9^, Parkinson’s disease^10,11^, rheumatoid arthritis^12^, and obesity and metabolic disorders induced by antipsychotic drugs^13^. However, AuNCs synthesized are usually complex mixtures, and the pharmacological studies of AuNCs mostly stayed on the level of disclosure of curative effects, yet the exact targets and their relationship with AuNCs molecular structures remained unclear and seldom discussed. It largely restricts the pharmaceutical development of AuNCs.

Autoimmune diseases (AIDs) are complex conditions characterized by the loss of immune tolerance to self-antigens and persistent autoreactive immune responses, which lead to the overproduction of autoantibodies and subsequent organ damages^14^. Until now, over 100 AIDs have been discovered, which affect more than 5% population in the world. The effective treatments for AIDs are still challenging and new drug discovery is much desirable^15^. Here we used two AuNCs with definite molecular structures, i.e., Au_18_(L-*N*- isobutyryl-L-cysteine)_14_ (Au_18_(L-NIBC)_14_) and Au_25_(L-NIBC)_18_, to study their different therapeutic effects in experimental autoimmune encephalomyelitis (EAE), a common-used animal model of multiple sclerosis (MS), and other AID animal models, and to identify the specific target in AID treatments.

Au_18_(L-NIBC)_14_ and Au_25_(L-NIBC)_18_ have the same ligand. Both of them possess cage structures composed of Au cores and staple structures comprising Au(Ⅰ) and ligands surrounding the Au cores^24,25^ (Fig. 1a and c). The two AuNCs samples were provided by Shenzhen Profound-View Pharmaceutical Technology Co., Ltd. Au_18_(L-NIBC)_14_ powder is dark green (inset in Fig. 1b) while that of Au_25_(L-NIBC)_18_ is brown (inset in Fig. 1d). Transmission electron microscopic (TEM) images indicated that both of them had uniform diameters of about 1 nm (Fig. 1b and d). Fig. 1e and f showed the chromatograms of C18 reversed-phase high-performance liquid chromatography (RP-HPLC) for the standard products of Au_18_(L-NIBC)_14_ and Au_25_(L-NIBC)_18_. The high-resolution mass spectra (HRMS) (Fig. 1g and h) of Au_18_(L-NIBC)_14_ and Au_25_(L-NIBC)_18_ were obtained in the m/z range of 1000-6000 Da by direct sampling using Thermo Q-Exactive Plus Hybrid Quadrupole-Orbitrap Mass Spectrometer. Both of them showed 3 main multi-charged ion peaks of z=4, 3, 2 and z=5, 4, 3, respectively. The corresponding subdivision spectra (black lines in insets of Fig. 1g and h) of the multi-charge ion peaks were highly consistent with theoretical calculations (red lines) from the molecular formulae. Other characterization data were shown in Extended Data Fig. 1.

**Fig. 1.**
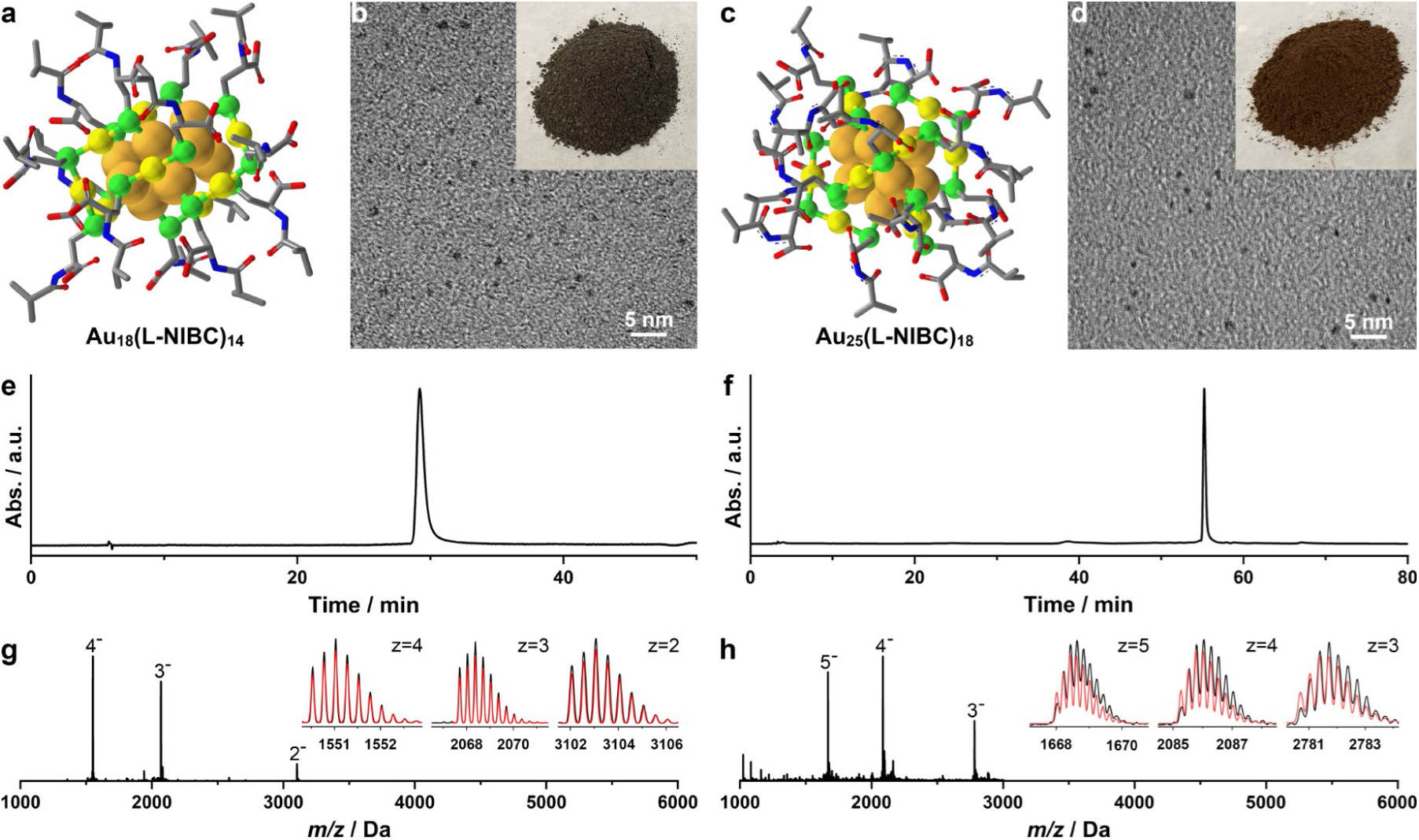
Characterization data of Au_18_(L-NIBC)_14_ sample. **a** and **c**, Schematic diagrams for cage structures of Au_18_(L-NIBC)_14_ (**a**) and Au_25_(L-NIBC)_18_ (**b**). Dark yellow: Au cores; light yellow: Au(Ⅰ); green: S; blue: N; red: O; grey: C. **b** and **d**, TEM images of Au_18_(L-NIBC)_14_ (**b**) and Au_25_(L-NIBC)_18_ (**d**). Insets show the photos of corresponding sample powders. **e** and **f**, C18 RP-HPLC chromatograms of Au_18_(L-NIBC)_14_ (**e**) and Au_25_(L-NIBC)_18_ (**f**). **g** and **h**, HRMS spectra for Au_18_(L-NIBC)_14_ (**g**) and Au_25_(L-NIBC)_18_ (**h**). Insets are subdivision spectra (black lines) of the multi-charge ion peaks and corresponding theoretical calculations results (red lines) from the molecular formulae.

Pilot study indicated that Au_18_(L-NIBC)_14_ had good therapeutic effect on motor behavioral dysfunctions of EAE mice, whereas the effect of Au_25_(L-NIBC)_18_ was poor. To obtain information about the potential mechanism and molecular target of Au_18_(L-NIBC)_14_ in EAE, we performed RNA sequencing (RNA-Seq) study on cortical tissues from Naive mice (Naive), EAE mice (EAE), and Au_18_(L-NIBC)_14_-treated EAE mice (Au_18_(L-NIBC)_14_). The Venn diagram (Extended Data Fig.2a) illustrated that there were 933 differentially expressed genes (DEGs) in Naive *vs.* EAE (comparison 1), which were all upregulated, and 270 DEGs in EAE *vs.* Au_18_(L-NIBC)_14_ (comparison 2), which were all downregulated. The overlapping region of two comparisons consisted of 222 DEGs, suggesting that they were involved in both EAE pathogenesis and therapeutic effects of Au_18_(L-NIBC)_14_.

We further conducted Kyoto Encyclopedia of Genes and Genomes (KEGG) pathway enrichment analysis of DEGs from comparison 1 and comparison 2 to figure out key pathways in EAE progression and the therapeutic effects of our drug. The top 20 enriched pathways for each comparison were presented in Extended Data Fig. 2b and 2c. Among them, the NOD-like receptor signaling pathway was the most significantly enriched in comparison 2, which ranked third in comparison 1. It suggested that this pathway played a crucial role in EAE progression and Au_18_(L-NIBC)_14_ intervention. The pathway diagrams displayed upregulated DEGs (red boxes) in comparison 1 (Extended Data Fig. 2d) and downregulated DEGs (green boxes) in comparison 2 (Extended Data Fig. 2e), which emphasized inflammasome-related molecular changes. We specially noted apoptosis-associated speck-like protein containing a CARD (ASC) because of its critical role in inflammasome activation within this pathway, which were significantly involved in both comparison 1 and comparison 2. Extended Data Fig. 2f illustrates the average gene expression levels of ASC. Compared to the Naive group, ASC expression was significantly elevated in the EAE group, however, treatment with Au_18_(L-NIBC)_14_ markedly reduced its expression.

ASC is a fibrous protein, which links activated inflammasome sensors to the effector pro-caspase-1 through oligomerization and subsequent fibrillation, resulting in the activation of caspase-1 and release of pro-inflammatory cytokines interleukin-1β (IL-1β) and IL-18, initiating or participating into a series of innate^19^ and adaptive immune-inflammatory cascades^20^. Without oligomerization and fibrillation, the inflammasome fails to be assembled, and downstream events will not happen^21,22^. Increasing evidences have also implicated that ASC plays an important role in the pathogenesis and progression of MS^16^ and other AIDs^17,18^. In previous works, we have reported that AuNCs possessed good anti-aggregation and fibrillation properties. Therefore, we speculated that ASC may be an important therapeutic target of Au_18_(L-NIBC)_14_ in EAE and other animal models of AIDs. This will be studied in detail in the following.

### *In vitro* study of anti-ASC oligomerization

The abilities of two AuNCs samples to inhibit ASC oligomerization were firstly studied *in vitro* by atom force microscopy (AFM) and dynamic light scattering (DLS). As shown in the AFM image (Fig. 2a), after 48 h of incubation at 37 ℃ and a neutral pH of 7.4, ASC alone strongly aggregated and large and thick fibrils were everywhere. However, when co-incubated with Au_18_(L-NIBC)_14_ of 10 ppm for 48 h, no obvious ASC fibrils could be observed (Fig. 2b). Co-incubation with Au_25_(L-NIBC)_18_ sample of 10 ppm could reduce the formation of ASC fibrils for a certain extent, but large ASC fibrils were still observed (Fig. 2c). DLS study (Fig. 2d) indicated that, when ASC was incubated alone, obvious ASC fibrillation (apparent sizes increased from several nm to hundreds of nm) occurred since about 25^th^ hour, before which the average apparent sizes of ASC maintained at 3-7 nm. Co-incubation with Au_25_(L-NIBC)_18_ largely postponed the starting time for ASC fibrosis to 49^th^ hour. However, when co-incubated with Au_18_(L-NIBC)_14_, ASC sizes still remained within 3-7 nm even after 53 hours of co-incubation. These data showed stronger inhibition effect of Au_18_(L-NIBC)_14_ than Au_25_(L-NIBC)_18_ on ASC fibrillation.

**Fig. 2.**
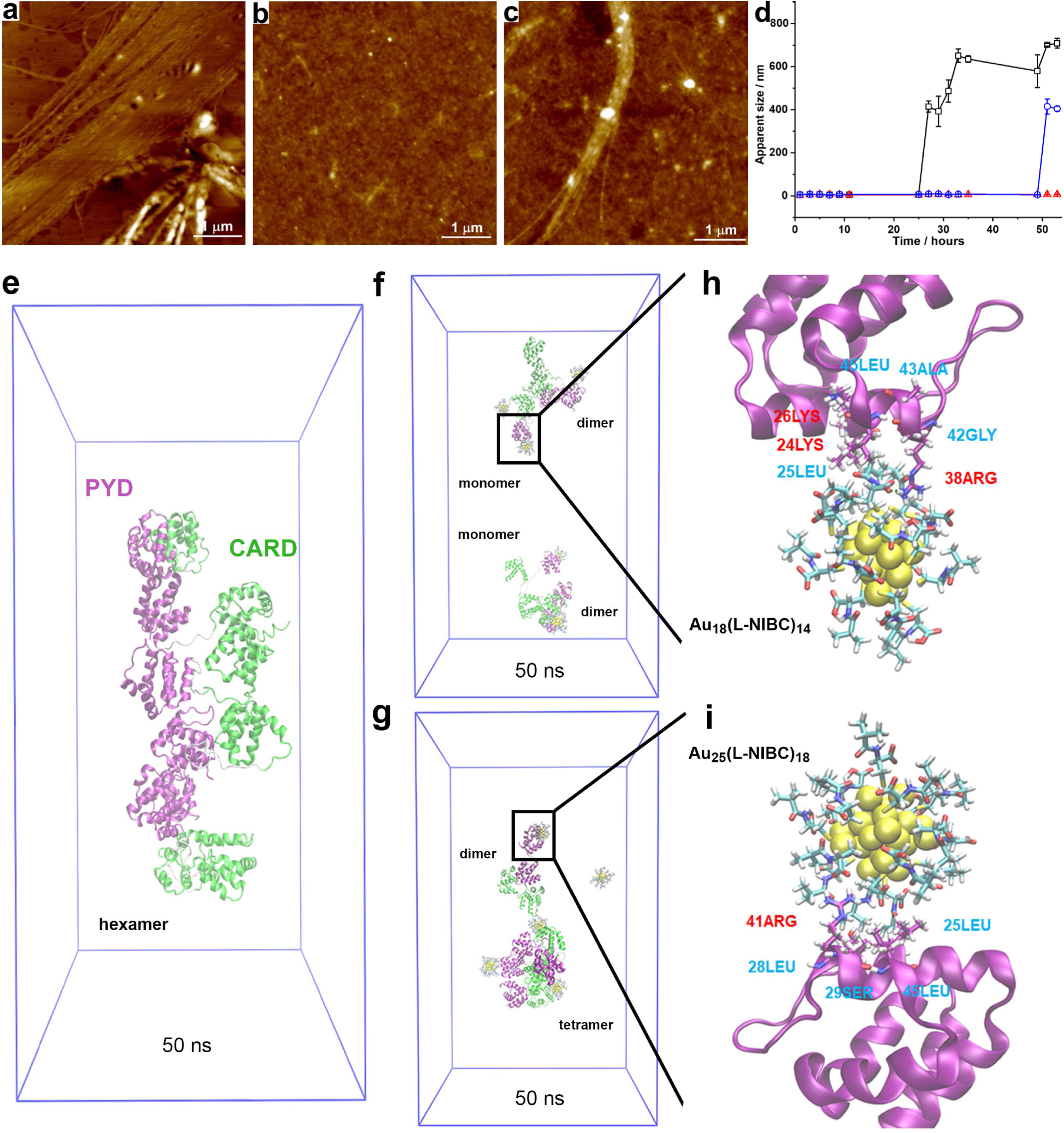
*In vitro* experiments show that Au_18_(L-NIBC)_14_ has stronger interaction with ASC than Au_25_(L-NIBC)_18_, and can effectively prevent ASC oligomerization. **a-c,** AFM images for ASC fibrillation after 48 hours of incubation without (**a**) or with the presence of Au_18_(L-NIBC)_14_ (**b**) or Au_25_(L-NIBC)_18_ (**c**). **d**, Apparent sizes of ASC measured by DLS experiment after incubation for different period of time without (black hollow squares) or with the presence of Au_18_(L-NIBC)_14_ (red hollow circles) or Au_25_(L-NIBC)_18_ (blue hollow triangles). **e-i**, Theoretical calculation results of all-atom MD simulation using 6 ASC molecules without (**e**) or with the presence of 6 Au_18_(L-NIBC)_14_ (**f**) or Au_25_(L-NIBC)_18_ (**g**), in which h and i are the corresponding magnification to show the electrostatic interactions (red) and hydrophobic interactions (light blue) between ASC and Au_18_(L-NIBC)_14_ (**h**) or Au_25_(L-NIBC)_18_ (**i**).

Theoretical calculation of all-atom molecular dynamics (MD) simulation using 6 ASC monomers indicated that ASC immediately aggregated to form hexamers within 5 ns via electrostatic and hydrophobic interactions between PYD domains, which is consistent with literatures^26,27^. The aggregates became obviously more compact when the calculation time was prolonged to 50 ns (Fig. 2e). The addition of 6 Au_18_(L-NIBC)_14_ into the same system effectively prevented the PYD domain of ASC to aggregate, and ASC could at most form dimers within 50 ns (Fig. 2f). However, with the presence of 6 Au_25_(L-NIBC)_18_, ASC still tend to aggregate into larger oligomers at 5 ns, and tetramers formed within 50 ns (**Fig. 2g**). MD simulation further reveals that both Au_18_(L-NIBC)_14_ and Au_25_(L-NIBC)_18_ can bind specifically to the PYD domains (but not CARD domains) of ASC. The difference is that Au_18_(L-NIBC)_14_ bound to the PYD domain via three pairs of electrostatic interactions between the negatively charged –COO^-^ of AuNCs and the positively charged 38ARG, 24LYS and 26LYS residues of PYD, which were strengthened by four pairs of hydrophobic interactions with 25LEU, 42GLY, 43ALA and 45LEU of PYD (**Fig. 2h**), whereas Au_25_(L-NIBC)_18_ bound to the PYD domain via only one pair of electrostatic interaction (41ARG) and 4 pairs of hydrophobic interactions (25LEU, 28LEU, 29SER, 45LEU) (**Fig. 2i**). It revealed stronger interaction between ASC and Au_18_(L-NIBC)_14_ than Au_25_(L-NIBC)_18_. These results are well consistent with the AFM and DLS experiments, indicating that Au_18_(L-NIBC)_14_ had a higher efficiency to inhibit ASC oligomerization than Au_25_(L-NIBC)_18_.

### Au_18_(L-NIBC)_14_ specifically targets ASC-dependent inflammasomes

NLR pyrin domain containing protein 3 (NLRP3) inflammasome is a typical and extensively studied ASC-dependent inflammasome^27^. The activation of NLRP3 and other inflammasomes inevitably leads to the maturation and release of pro-inflammatory cytokines interleukin (IL)-1β and IL-18^28^. We firstly evaluated the abilities of Au_18_(L-NIBC)_14_ and Au_25_(L-NIBC)_18_ (concentration: 20 ppm) to inhibit NLRP3 inflammasome activation in cell level^29^. As shown in Fig. 3a and b, NLRP3 inflammasome activation resulted in great increase of IL-1β and IL-18 levels. The addition of Au_18_(L-NIBC)_14_ significantly reduced IL-1β and IL-18 levels, respectively. However, for Au_25_(L-NIBC)_18_, although the reduction for IL-1β and IL-18 levels were also remarkable, the reduction extents are much lower than Au_18_(L-NIBC)_14_.

**Fig. 3.**
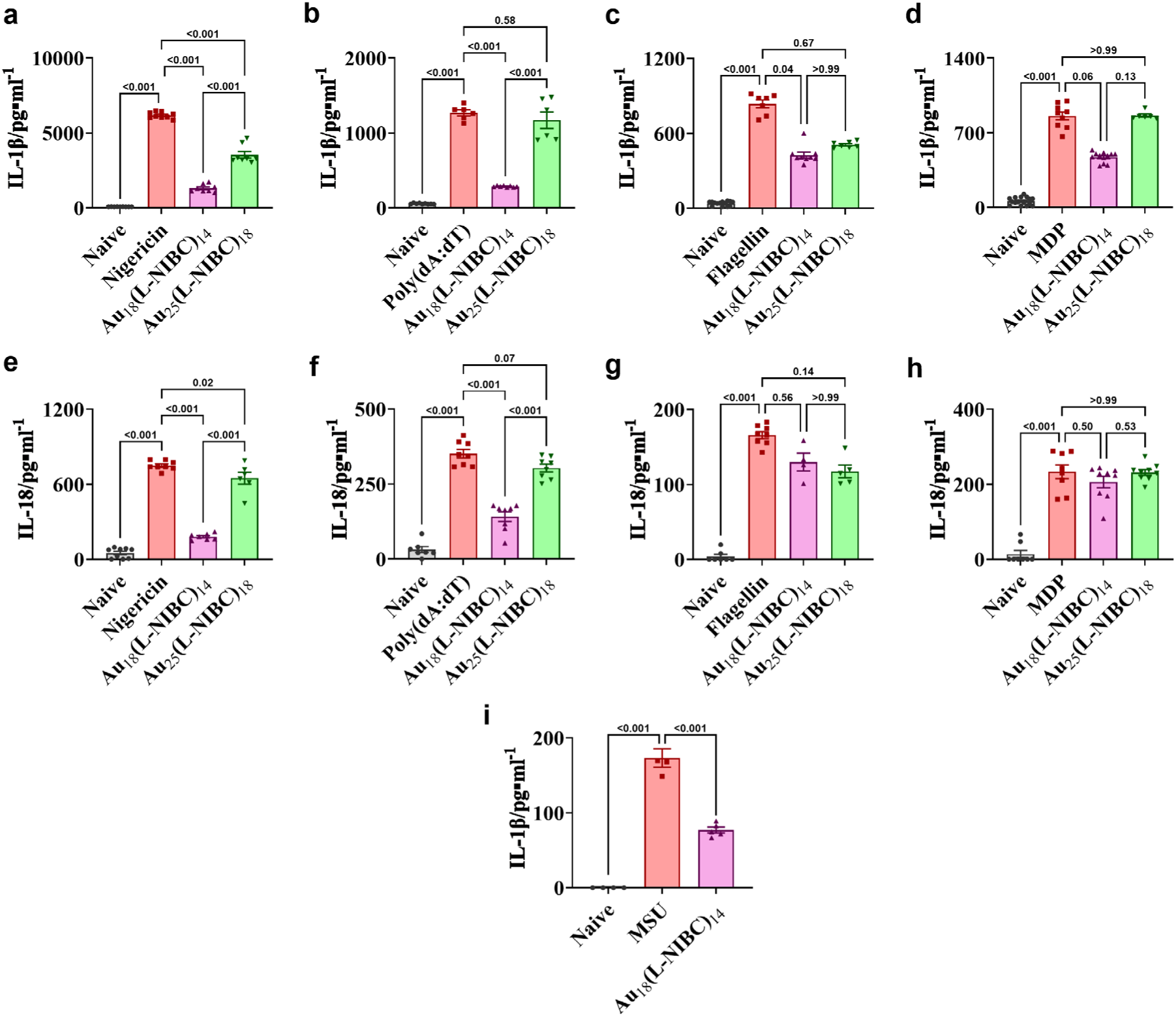
Cell experiments (a-h) show that Au_18_(L-NIBC)_14_ inhibit the activation of ASC-dependent inflammasomes specifically. PMA differentiated THP-1 cells stimulated by LPS plus nigericin (**a**, **e**), dA:dT (**b**, **f**), flagellin (**c**, **g**), and MDP (**d**, **h**) generate NLRP3, AIM2, NLRC4, and NLRP1 inflammasomes, respectively. **a-d**, production of IL-1β measured by ELISA with or without Au_18_(L-NIBC)_14_ treatment, in comparison with Au_25_(L-NIBC)_18_ treatment. **e-h**, corresponding results of IL-18. Experiment of an NLRP3-dependent murine peritonitis model shows that the inhibition effects of Au_18_(L-NIBC)_14_ on inflammasome activation also works *in vivo*. **i**, IL-1β concentration in the peritoneal cavity of C57BL/6 mice injected with MSU crystals with or without Au_18_(L-NIBC)_14_ treatment.

We then studied the effects of our drugs on another ASC-dependent inflammasome— AIM2 (absent in melanoma 2)^30^ and two ASC-independent inflammasomes—NOD-like receptor CARD domain containing protein 4 (NLRC4)^31^ and NOD-like receptor pyrin domain containing protein 1 (NLRP1)^32^. As shown in Fig. 3b-d and Fig.3f-h, after stimulation for three inflammasomes, IL-1β levels and IL-18 levels all significantly increased. Au_18_(L-NIBC)_14_ strongly inhibited AIM2 activation (Fig. 3b,f). But for NLRC4 and NLRP1, although Au_18_(L-NIBC)_14_ reduced IL-1β for some extents (Fig. 3c,d), evidential decreases were not observed for IL-18 (Fig. 3g,h). Considering that different inflammasomes may interact with each other and thus ASC may indirectly influence NLPC4 and NLRP1, we conclude that Au_18_(L-NIBC)_14_ could hardly inhibit the activation of NLPC4 and NLRP1 directly. On the other hand, Au_25_(L-NIBC)_18_ sample did not show any evidential inhibition effects on all three inflammasomes (Fig. 3b-d and f-h).

These results revealed that Au_18_(L-NIBC)_14_ could strongly and specifically inhibit the activation of ASC-dependent inflammasomes in cell level. But for Au_25_(L-NIBC)_18_, the inhibition effect is relatively very weak, which is also lack of specificity. An NLRP3 dependent murine peritonitis model^33^ further indicated that Au_18_(L-NIBC)_14_ could inhibit ASC-dependent inflammasome activation *in vivo*, in which IL-1β level was significantly reduced (Fig. 3i).

### Therapeutic effects on EAE mice

Motor impairment is one of the most important symptoms of MS and EAE. MCC950 is a small molecular inhibitor, which can specifically inhibit NLRP3 inflammasome activation via interaction with the Walker B motif within the NLRP3 NACHT domain^34^. It has been extensively studied due to its good therapeutic effects in NLRP3-associated syndromes including those of EAE^35^. AFM and DLS experiments indicated that MCC950 could not inhibit ASC oligomerization, and cell experiments showed that it could effectively inhibit NLRP3 activation but ineffective to other inflammasomes, e.g., AIM2, NLRC4 and NLRP1 (Extended Data Fig. 3). Adrenocortical hormone prednisone is a powerful immunosuppressive agent which has been extensively used in MS treatment. Here MCC950 and prednisone were used as the positive controls for the study of motor dysfunctions in EAE experiments.

During the EAE experiment, the clinical scores of mice were recorded every day to study the motor behavioral changes of animals. As expected, in the EAE group, clinical symptoms such as limp tail, appeared within two weeks post EAE induction, their hind limb became dragging or complete paralysis at the end of the third week. EAE mice treated with MCC950 (administration routes were all intraperitoneal injection unless otherwise specified) started to exhibit early clinical syndromes within 2 weeks, serious motor dysfunction, e.g., hind limb paralysis, rarely happened until the experiment finished, but mild symptoms such as limp tail and hind limb inhibition were still very common (Fig. 4a), which is similar to literatures^35^. EAE mice with prednisone treatment also started to exhibit early clinical syndromes within 2 weeks, and the clinical score apparently decreased since day22 from 1.30 ± 0.21 to 0.20 ± 0.13 on day30 (Fig. 4b). It indicated that both MCC950 and prednisone had good effects to ameliorate the motor impairment of EAE mice. Interestingly, EAE mice administrated with Au_18_(L-NIBC)_14_ (20 mg/kg) did not show any symptom, and almost all the clinical scores maintained 0 during the whole process of experiment (Fig. 4c). In contrast, the effect of Au_25_(L-NIBC)_18_ administration with the same dosage was very weak (Fig. 4c). Dosage screening experiments of Au_18_(L-NIBC)_14_ (Fig. 4d) showed that dosages of 1 mg/kg and above could all give the complete inhibition effect on clinical scores, indicating that the effective dosage should be equal to or lower than 1 mg/kg. Oral administration also gave good result on clinical scores (Fig. 4e). These data show that Au_18_(L-NIBC)_14_ possesses a very good therapeutic effect on motor dysfunctions of EAE mice, which is superior to that of MCC950 and prednisone.

**Fig. 4.**
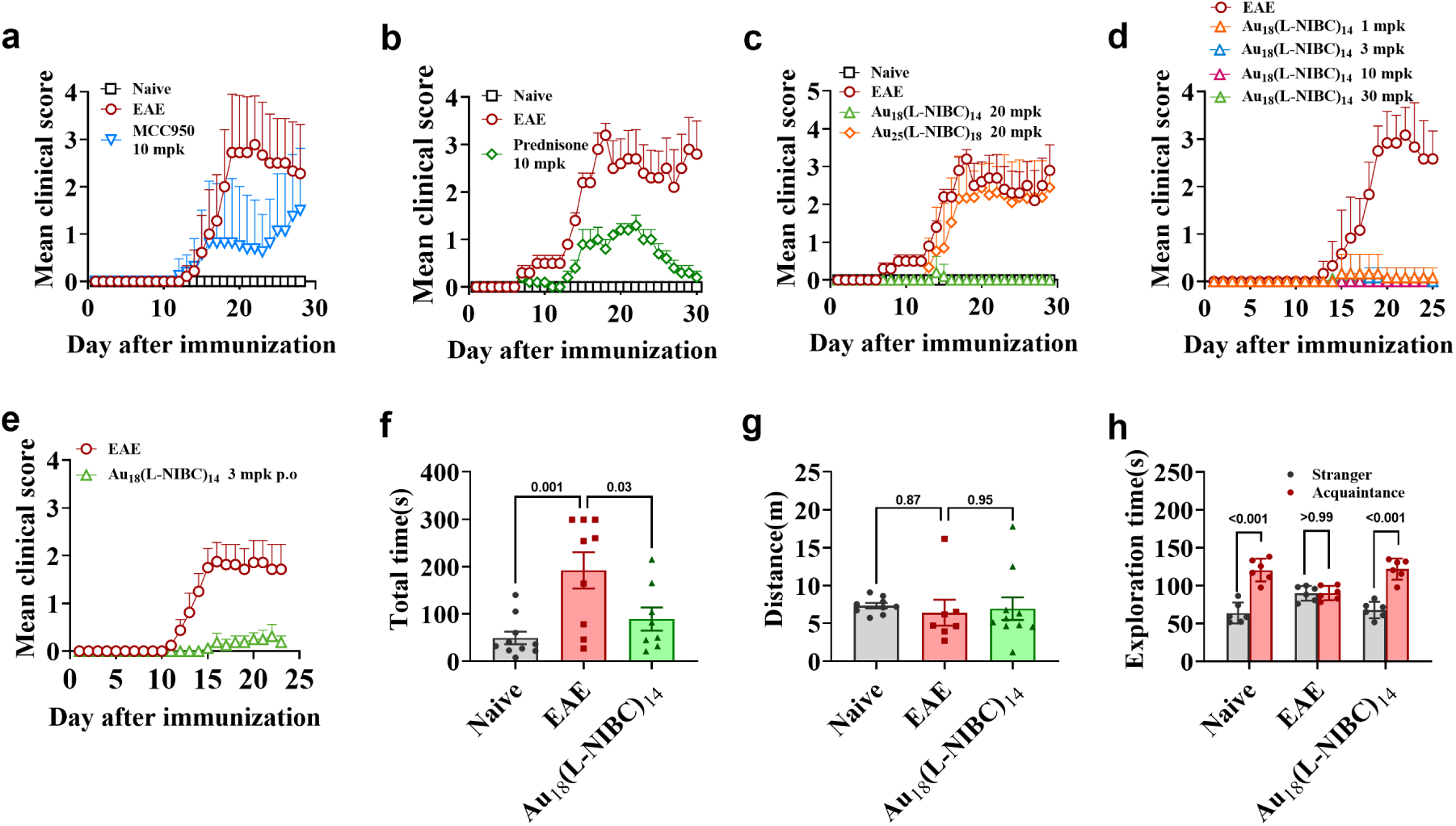
Au_18_(L-NIBC)_14_ exhibited excellent capacity to eliminate the motor dysfunction (a-e) and early memory and cognitive impairment of EAE mice (f-h). **a-e**, clinical scores of motor dysfunction. **a**, MCC950 treated EAE mice. **b**, prednisone treated EAE mice. **c**, comparison between EAE mice treated with Au_18_(L-NIBC)_14_ and Au_25_(L-NIBC)_18_, respectively. **d**, EAE mice treated with different dosages of Au_18_(L-NIBC)_14_. **e**, EAE mice with oral administration (p.o) of Au_18_(L-NIBC)_14_. **f**, the travel time for Au_18_(L-NIBC)_14_ treated EAE mice to find the fixed hole of escape compartment in Barnes Maze test. **g**, the travel distances for Au_18_(L-NIBC)_14_ treated EAE mice during 300-second period in open field test. **h**, the exploration time for Au_18_(L-NIBC)_14_ treated EAE mice on a “stranger” mouse and an “acquaintance” mouse in the three-chambered social interaction test.

Impairment of memory and cognition is another common and devasting manifestation of MS. Literatures have shown that in EAE mice, memory and cognitive impairment has already occurred before motor dysfunction^36,37^. We used Barnes maze test to evaluate the spatial memory of mice, and three-chamber sociability test to assess the cognition of mice. In Barnes maze test, mice in EAE group spent significantly longer time to find the fixed hole of escape compartment during the 300-second period of test, compared with those in the Naive group, indicating an obvious impairment in spatial memory. Au_18_(L-NIBC)_14_ treatment greatly decreased the time for EAE mice to find the escape hole compared with the EAE model group (Fig. 4f). In three-chamber sociability test, EAE mice did not show evidential motor dysfunction in open field experiment (Fig. 4g), which guarantees that the results of sociability test were not influenced by the motor behavioral ability. In this test, mice in Naive group displayed normal social memory and spent more time exploring the “stranger” mouse than the “acquaintance” mouse (Fig. 4h). However, mice in EAE group showed no preference for either “stranger” mouse or “acquaintance” mouse, indicating the impairment of cognition. Au_18_(L-NIBC)_14_ treated EAE mice also exhibited distinct preference for “stranger” mouse rather than “acquaintance” mouse, demonstrating that Au_18_(L-NIBC)_14_ treatment effectively prevented cognitive impairment of EAE mice.

The effect of Au_18_(L-NIBC)_14_ on ASC-dependent inflammasome activation was further investigated in spinal cords of EAE mice at the end of the EAE experiment (day30). IL-1β, IL-18 and caspase-1 are the essential products of inflammasome activation. Enzyme-linked immunosorbent assay (ELISA) indicated that IL-1β level increased significantly in EAE group compared to the Naive group (Fig. 5a). Remarkably, treatment with Au_18_(L-NIBC)_14_ completely suppressed the IL-1β level to the Naive level, in which the effect was obviously better than that of MCC950 (p=0.01) (Fig. 5a). Similarly, Au_18_(L-NIBC)_14_ also significantly reduced the levels of IL-18 (Fig. 5b) and caspase-1 (Fig. 5c). In contrast, the effect of Au_25_(L-NIBC)_18_ was very weak for IL-1β, IL-18 and caspase-1. ASC speck formation is a hallmark of ASC-dependent inflammasome activation^38^. It was further assessed using immunofluorescence colocalization, in which ASC specks were stained red, while the nuclei stained with DAPI were blue. Statistic results showed that ASC speck ratio in spinal cords of EAE mice significantly increased compared to the Naive group. Au_18_(L-NIBC)_14_ treatment reduced the ASC speck ratio to almost the same level as the Naive group (Fig. 5d,e). However, Au_25_(L-NIBC)_18_ did not show the effect evidentially (Fig. 5d,e). These results indicated that Au_18_(L-NIBC)_14_ could effectively inhibit the inflammasome activation in the CNS of EAE mice, but Au_25_(L-NIBC)_18_ could not.

**Fig. 5.**
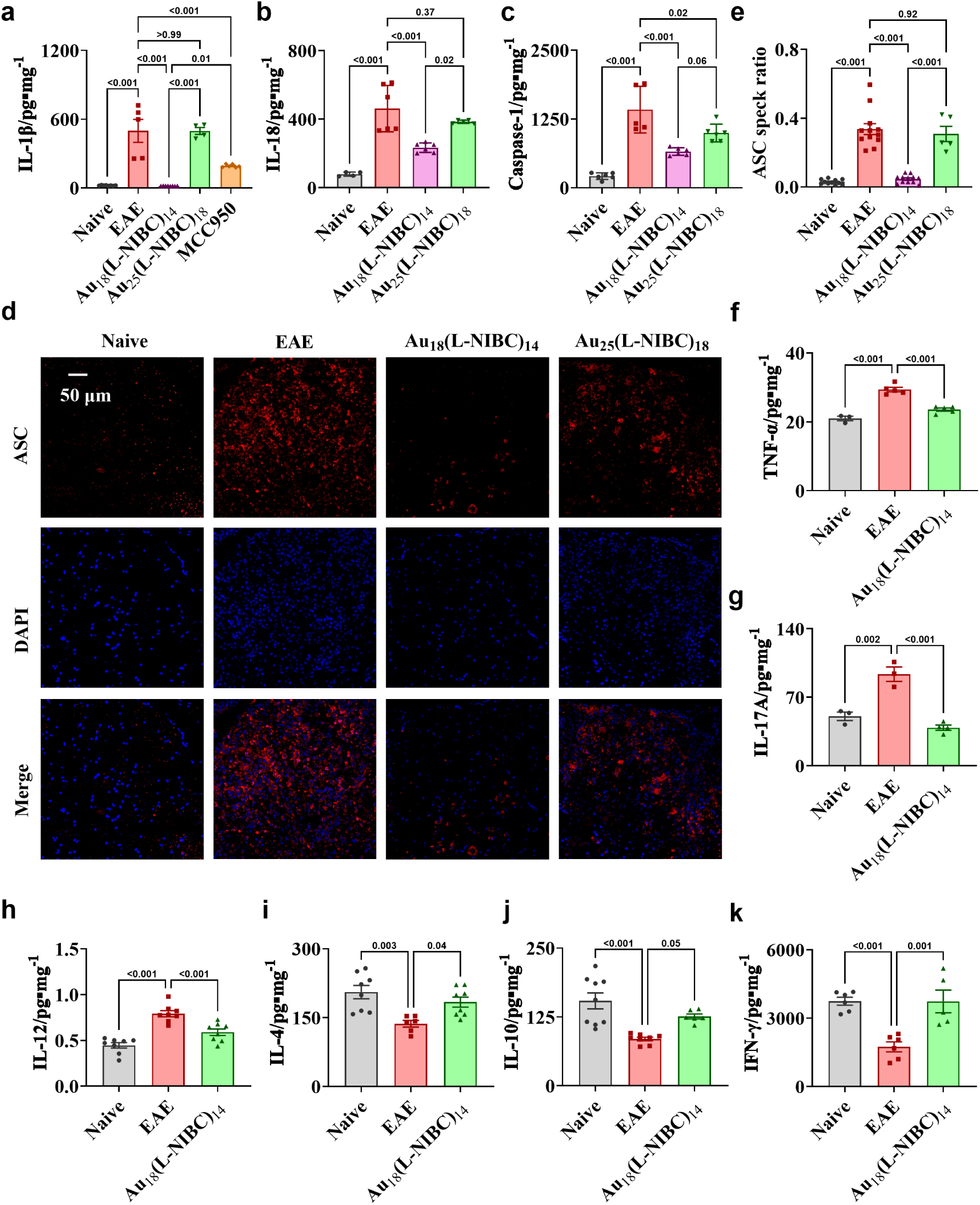
Au_18_(L-NIBC)_14_ demonstrates robust inhibition of CNS inflammation in EAE mice. **a-c**, ELISA results show that Au_18_(L-NIBC)_14_ significantly reduced the levels of IL-1β (**a**), IL-18 (**b**), and caspase-1 (**c**) in the spinal cords of EAE mice, while Au_25_(L-NIBC)_18_ showed no evidential inhibitory effects. **d-e**, Representative fluorescence microscopy images (**d**) and quantitative analysis (**e**) reveal a marked reduction in ASC specks in the spinal cords of EAE mice treated with Au_18_(L-NIBC)_14_, indicating effective inhibition of inflammasome activation. Au_25_(L-NIBC)_18_ did not reduce the ratio of ASC specks. **f-k**. Regulation of other cytokines: pro-inflammatory cytokines TNF-α (**f**), IL- 17A (**g**) and IL-12 (**h**) were significantly reduced by Au_18_(L-NIBC)_14_; while anti-inflammatory cytokines IL-4 (**i**), IL-10 (**j**) and IFN-γ (**k**) levels were significantly increased by Au_18_(L-NIBC)_14_.

The inflammasome activation and the release of IL-1β, IL-18, and caspase-1 will further activate or regulate innate immune cells, e.g. macrophages and microglia^39^, and the adaptive immune cells, e.g. T cells^20^ and B cells^40,41^. Therefore, we next studied the effects of Au_18_(L-NIBC)_14_ on the production of other cytokines in spinal cords of EAE mice. As shown in Fig. 5f-h, pro-inflammatory cytokines TNF-α, IL-17A, and IL-12 were markedly reduced by Au_18_(L-NIBC)_14_ treatment. Conversely, Au_18_(L-NIBC)_14_ significantly increased the levels of anti-inflammatory cytokines IL-4, IL-10 and IFN-γ (Fig. 5i-k), which were substantially suppressed by EAE modelling. We also measured the expression of several other cytokines, e.g. IL-6, IL-23, IL-15, IL-35 and GM-CSF, but we did not observe any statistical difference among the Naive group, EAE group, and the Au_18_(L-NIBC)_14_ treated group (*p* all larger than 0.05). These results revealed that Au_18_(L-NIBC)_14_ possessed a powerful capacity of anti-CNS inflammation, which can comprehensively restore the cytokine homeostasis in the CNS of EAE mice.

The infiltration of immune cells into the CNS from the peripheral and demyelination of axons are important pathological hallmarks of MS/EAE^42^. Fig. 6a and b show typical H&E staining image and the statistic results of the spinal cords of mice on day21 after EAE induction, in which the nuclei of the infiltrated immune cells under light microscopy were blue. In EAE group, the number of immune cells infiltrating into spinal cords significantly increased compared to the Naive group. Au_18_(L-NIBC)_14_ treated EAE mice strongly decreased the immune cell infiltration compared to the EAE group. However, in EAE mice treated with Au_25_(L-NIBC)_18_, the reduction of immune cell infiltration was not obvious compared to the EAE group. It indicated that Au_18_(L-NIBC)_14_ strongly inhibited immune cells infiltration into CNS from the peripheral, but the effect of Au_25_(L-NIBC)_18_ was weak.

**Fig. 6.**
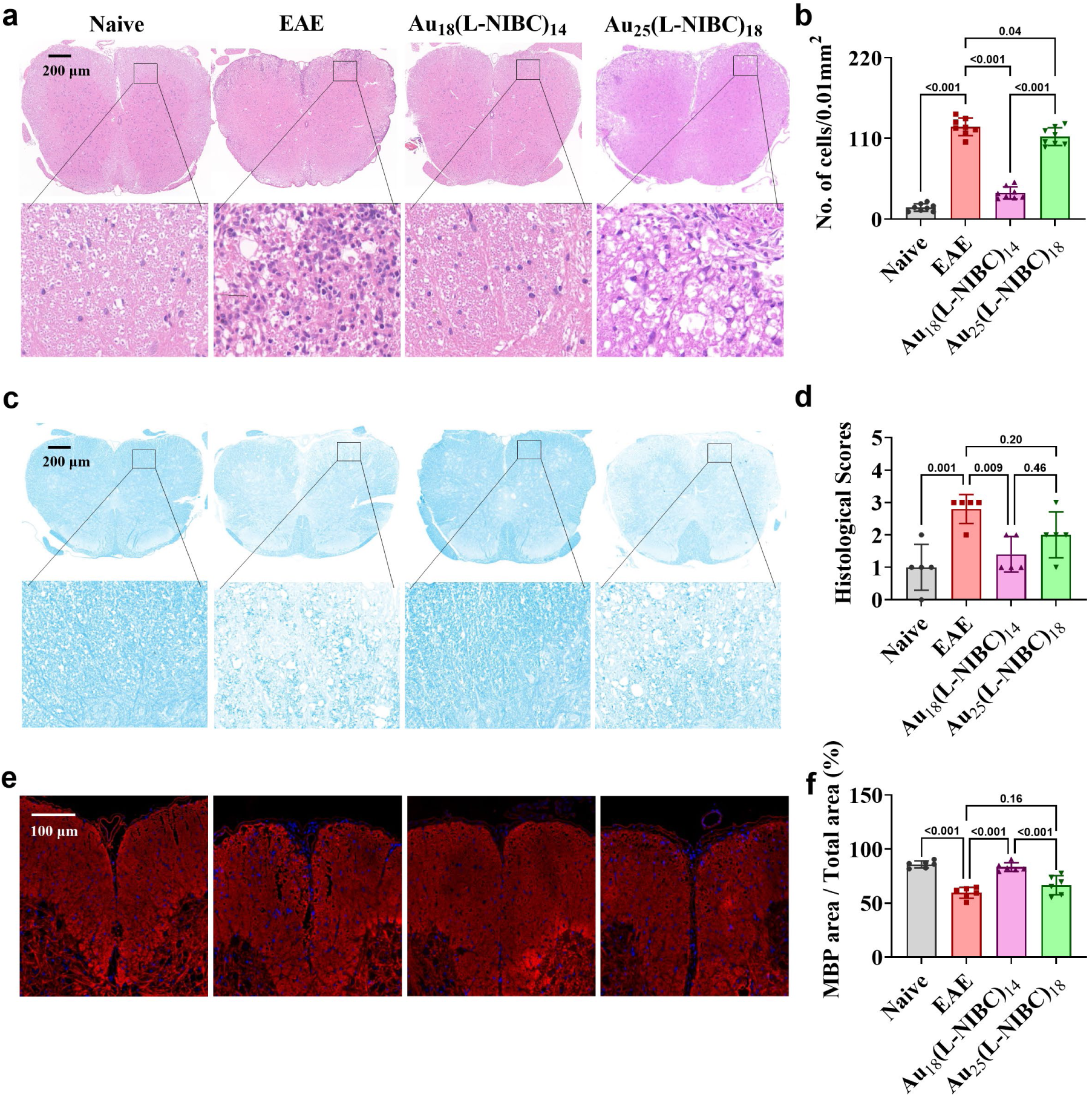
Au_18_(L-NIBC)_14_ shows excellent effect to inhibit immune cells infiltration into CNS from peripheral and prevent axon demyelination. **a**, representative H&E staining pictures of spinal cords for mice in Naive group, EAE group, Au_18_(L-NIBC)_14_ treated group, and Au_25_(L-NIBC)_18_ treated group, respectively. The nuclei of the infiltrated immune cells were stained to blue under light microscopy. **b**, corresponding statistic results. **c**, representative LFB staining pictures of spinal cords for mice in Naive group, EAE group, Au_18_(L-NIBC)_14_ treated group, and Au_25_(L-NIBC)_18_ treated group, respectively. **d**, corresponding statistic results for histological scores. **e**, representative MBP staining pictures of spinal cords for mice in Naive group, EAE group, Au_18_(L-NIBC)_14_ treated group, and Au_25_(L-NIBC)_18_ treated group, respectively, in which MBP was stained to red. **f**, corresponding statistic results for the percentages of MBP stained areas in the white matter of spinal cords.

Next, demyelination of axons was assessed in spinal cord of mice using both Luxol fast blue (LFB) staining and myelin basic protein (MBP) staining. In LFB staining, myelin is visualized and stained blue under light microscopy, in which the demyelination extent of axons can be scored based on the detection of focal white matter areas lacking LFB staining using the relative three-point scale. As shown in Fig. 6c and Fig. 6d for the LFB staining results, in spinal cords of mice in Naive group, only small areas of demyelination were observed (histological score: 1.00 ± 0.71). The mice in EAE group showed high histological score of 2.80 ± 0.20, indicating severe demyelination of axons in spinal cords. Au_18_(L-NIBC)_14_ treatment significantly reduced the histological score to 1.40 ± 0.24, whereas Au_25_(L-NIBC)_18_ treatment did not show evidential reduction compared to the EAE group. The MBP staining experiment gave the similar results, as shown in Fig. 6e and Fig. 6f. Both results demonstrated that Au_18_(L-NIBC)_14_ could effectively prevent demyelination of axons, but effect of Au_25_(L-NIBC)_18_ was weak.

The BBB is an active and selective barrier that shields the brain from exogenous insults. BBB disruption is also an important hallmark in MS/EAE^43,44^, which is widely involved in the pathogenesis and disease progression including infiltration of immune cells into CNS, *etc*. Here we used FITC-Dextran (10kDa) staining method and Evans’s blue (EB) dye extravasation method to study the effects of our drugs on BBB protection. FITC-Dextran and EB cannot pass through the integral BBB but can penetrate into the CNS when the BBB is disrupted. Extended Data Fig. 4a showed the representative images of EB dye extravasation in the brains and spinal cords. Examination of cerebral cortices and spinal cords revealed obvious blue staining in mice of EAE group and Au_25_(L-NIBC)_18_ treated EAE mice, which was not present in either Naive or Au_18_(L-NIBC)_14_ treatment EAE mice. It indicated that EAE modeling destructed the BBB integrity, whereas Au_18_(L-NIBC)_14_ treatment effectively prevented BBB destruction. Quantitative analysis (Extended Data Fig. 4b,c) showed significantly higher content of EB signals in both cerebral cortices and spinal cords of EAE group compared to the Naive group. Au_18_(L-NIBC)_14_ greatly reduced EB contents in both cerebral cortices and spinal cords to nearly the level of the Naive group. Au_25_(L-NIBC)_18_ also reduced the EB contents in cerebral cortices and spinal cords, but the reduction extents were significantly smaller than Au_18_(L-NIBC)_14_. The quantitative data of FITC-Dextran staining experiment (Extended Data Fig. 4d,e) gave similar results, and the effect was even more distinct.

Extended Data Fig. 4f and g displayed the Western-Blot (WB) results for occludin, a typical tight junctional protein, in brain capillary isolated from cerebral cortices and spinal cords of mice. In EAE group, occludin expression in both cerebral cortices and spinal cords decreased significantly compared to the Naive group. Au_18_(L-NIBC)_14_ treatment effectively prevented the decrease of occludin expression in cerebral cortices and spinal cords, but Au_25_(L-NIBC)_18_ treatment did not show evidential effect in this aspect (Extended Data Fig. 4h,i). These results demonstrated an excellent BBB protection property of Au_18_(L-NIBC)_14_.

Overall, Au_18_(L-NIBC)_14_ exhibits excellent therapeutic effects toward EAE mice, which include: complete inhibition of motor dysfunctions and relief of early-cognitive impairments, comprehensive regulation and restoration of abnormal pro- and anti-inflammatory cytokines in the CNS, prevention of immune cell infiltration from peripheral to the CNS and axons demyelination, and BBB protection.

### Therapeutic effects in other AID animal models

Numerous inflammatory cytokines have been identified as efficacious targets in drug development for various AIDs. For instance, drugs respectively targeting IL-1β, IL-6, IL- 12, or IFN-γ show promise in the treatment of Inflammatory Bowel Disease (IBD), and IL-17A and IL-23 are effective targets for psoriasis drug development. On the other hand, IBD^45,46^, psoriasis^47^, SLE^48^, and many other AIDs^49^ were reported to be highly relevant to varieties of inflammasomes. The inhibitory effects on multiple ASC-dependent inflammasomes and powerful capacity to restore cytokine homeostasis of Au_18_(L-NIBC)_14_ inspire us to study its potential in the treatments of other AIDs.

IBD is a chronic, multifactorial disorder that affects the gastrointestinal tract, particularly the colon and small intestine. It is primarily classified into two types: Crohn’s disease (CD) and ulcerative colitis (UC). To assess the therapeutic potential of Au_18_(L-NIBC)_14_ in IBD, we employed 2,4,6-trinitrobenzenesulfonic acid (TNBS)-induced, oxazolone (OXZ)-induced and Dextran sulfate (DSS)-induced colitis mice models to simulate CD and UC, respectively.

In the TNBS-induced colitis model (Extended Data Fig. 5a-d), mice exhibited significant weight loss, increased DAI scores, shortened colon length and increased colon ratio (the ratio of colon weight to colon length). Au_18_(L-NIBC)_14_ treatment at concentrations of 3 mg/kg and 6 mg/kg significantly increased the weight of mice, decreased the DAI score, increased colon length and decreased colon ratio compared with the mice in model group, which exhibited an apparent dosage-dependent result. The detection of inflammatory cytokines (Extended Data Fig. 5e-g) indicated that Au_18_(L-NIBC)_14_ could significantly inhibit the elevation of IL-1β, IL-6 and TNF-α induced by TNBS modeling in a dosage-dependent manner. Extended Data Fig. 5h and i showed the representative H&E staining results of colon slices and the corresponding pathological scores, indicating that Au_18_(L-NIBC)_14_ could efficiently ameliorate the pathological damage of colon. The DSS-induced colitis model (Extended Data Fig. 6a-d) and OXZ-induced colitis models (Extended Data Fig. 6e-h) gave similar results. These results revealed an excellent curative prospect in IBD treatment.

IMQ-induced psoriasis mouse model (Extended Data Fig. 7 was used to evaluate the potential of Au_18_(L-NIBC)_14_ in psoriasis treatment. IMQ treated model mice exhibited typical psoriasis-like inflammatory responses on the back and increased ear thickness compared to the Naive group. Au_18_(L-NIBC)_14_ treatment at concentrations of 3 mg/kg and 6 mg/kg significantly reduced the PASI scores on the back of mice and the ear thickness increase, compared to the untreated model mice in vehicle group. The data showed that Au_18_(L-NIBC)_14_ was effective in IMQ-induced psoriasis mouse model.

Female MRL/MpJ-Faslpr mouse model was used to study the therapeutic effects of Au_18_(L-NIBC)_14_ on systemic lupus erythematosus (SLE) (Extended Data Fig. 8. Au_18_(L-NIBC)_14_ treatment greatly reduced the urine albumin, the ratio of urine albumin to urine creatinine (ACR), and the blood urea nitrogen (BUN) of MRL/MpJ-Faslpr mice (Extended Data Fig. 8a-c), indicating effective amelioration of renal dysfunction. Extended Fig. 8d-g showed that Au_18_(L-NIBC)_14_ could significantly decrease the extent of renal fibrosis and glomerulosclerosis, indicating an excellent renal protection effect.

## Discussion and conclusions

Above results indicate that Au_18_(L-NIBC)_14_ can act as a promising candidate for drug discovery of MS and other AIDs.

First, excellent therapeutic effects of Au_18_(L-NIBC)_14_ on EAE model including extensive restoration of CNS cytokine homeostasis, inhibition of immune cell infiltration from the peripheral to the CNS, prevention of neuron axon demyelination, BBB protection and complete prevention of motor dysfunctions and relief of early memory and cognitive impairment, point out a bright prospect in MS treatment.

Second, the significantly better pharmaceutical effects compared to MCC950 and prednisone demonstrate the target superiority of ASC oligomerization. Blocking ASC oligomerization prevented the activation of multiple ASC-dependent inflammasomes, which apparently surpassed MCC950 (only targeting NLRP3) in mechanism. As a result, Au_18_(L-NIBC)_14_ administration not only resulted in complete inhibition of IL-1β to normal level (MCC950 could only partially inhibit the IL-1β level), but also showed powerful capacity to restore the homeostasis of other pro- and anti-inflammation cytokines in the CNS of EAE mice. This is distinctly superior to conventional immunosuppressants which mostly target one cytokine. More importantly, the generation and development of EAE involve multiple inflammasomes^50^, which further display the target advantages of ASC oligomerization in MS drug discovery.

Third, significant therapeutic effects in animal models of other AIDs, including IBD, psoriasis and SLE, illustrate good extensibility of Au_18_(L-NIBC)_14_ in treatments of different AID indications. This also support the superiority of targeting ASC oligomerization in AID drug discovery.

Fourth, well-defined molecular structure, specific target and clear mechanism, and excellent pharmacological activities, figure out a feasible route to solve the problems of druggability for Au_18_(L-NIBC)_14_ and other AuNCs. They form an important basis for preclinical studies and investigational new drug (IND) application, e.g., quality specification, pilot production, pharmacokinetics, pharmacodynamics and safety pharmacology.

## Methods

### Materials

Au_18_(L-NIBC)_14_, Au_25_(L-NIBC)_18_ samples were provided by Shenzhen Profound-View Pharmaceutical Technology Co., Ltd.. Ultrapure water with resistivity of 18.2 MΩ·cm was prepared by Milli-Q. All other reagents and antibodies were purchased from commercial vendors. Methanol (HPLC, Thermo Fisher); NaOH (AR, Sinopharm), Ltd.; KH_2_PO_4_ (AR, Sinopharm); NaH_2_PO_4_ (AR, Sinopharm); Phorbol 12-myristate 13-acetate (HY-18739, MedChemExpress); Nigericin (HY-100381, MedChemExpress); poly(dA:dT) (tlrl-patn, InvivoGen); Flagellin (SRP8029, Sigma); MDP (M9508, Sigma); Lipofectamine^TM^ 2000 Transfection Reagent (11668019, Thermo Fisher); Human IL-1β ELISA Kit (RK00001, ABclonal); Human IL-18 ELISA Kit(RK00176, ABclonal); Mouse IL-1β ELISA Kit (RK00006, ABclonal); Mouse IL-18 ELISA Kit (BMS618-3, Invitrogen); Mouse TNF-α ELISA Kit (RK00027, ABclonal); Mouse Caspase-1 ELISA Kit (AG-45B-0002-KI01, AdipoGen); Mouse IL-17A ELISA Kit (RK00039, ABclonal); Mouse IFN-γ ELISA Kit (RK00019, ABclonal); Mouse IL-4 ELISA Kit (RK00036, ABclonal); Mouse IL-10 ELISA Kit (RK00016, ABclonal); Mouse IL-12 Elisa Kit (E- EL-M3062, Elabscience); Hematoxylin and Eosin Staining Kit (C0105S, Beyotime); Luxol Fast Blue Stain Solution (G3242, Solarbio); BCA Protein Assay Kit (P0012, Beyotime); RIPA Lysis Buffer (P0013B, Beyotime); ECL (S6009L, US Everbright); Gota Serum (16210064, Thermo Fisher); Antifade Mounting Medium with DAPI (A4084, US Everbright); Rabbit anti-MBP antibody (78896T, Cell Signaling Technology); Rabbit anti-ASC antibody (67824, Cell Signaling Technology); Mouse anti-Occludin antibody (33-1500, Invitrogen); Mouse anti-GAPDH antibody (MA1- 16757, Invitrogen); Goat anti-mouse antibody (A0216, Beyotime); Alexa Fluor 568 (A- 11011, Thermo Fisher).

### Characterizations for AuNCs

The TEM images of AuNCs were performed on a JEOL 200CX TEM (JEOL Ltd., Tokyo, Japan) at 200 kV. For preparation of the TEM sample, a diluted solution of AuNCs in ethanol/ultrapure water was dropped on a carbon coated TEM grid (Cu, 300 mesh, Electron Microscopy Sciences, Hatfield, PA) and then dried in air. UV spectra were recorded on Shimadzu UV-1900 UV-vis spectrophotometer with a range of 300 to 900 nm at a scan rate of 0.5 nm s^‒1^. SEC chromatography was conducted using an Thermo UltiMate 3000 HPLC system equipped with an autosampler. In this study, a polymer-based columns (OHpak SB-802.5 HQ, 300 mm × 8 mm) was used, and the detection was made with a DAD detector at 254 nm. AuNCs dissolved in ultrapure water was sampled by injection (10 μL), and then eluted by the mobile phase (0.1 M phosphate buffer, pH 7.00) at a flow rate of 0.4 ml‧min^-1^ for 50 min. HPLC analysis was performed on a Shimadzu system (Prominence-i LC-2030c 3D plus) with PDA detector (detection wavelength: 254 nm). The reversed phase column (Agilent ZORBAX Eclipse Plus C18, 250 mm × 4.6 mm) was used at a constant temperature of 30 ℃. Samples were separated by gradient elution at a flow rate of 0.8 ml‧min^-1^ with mobile phase composed of 50 mmol/L KH_2_PO_4_ (pH6.8) (eluent A) and methanol (eluent B) and a gradient program: 22% B for 40 min, 22%-60% B gradiently for 30 min, 60% B for 10 min, 90% B for 10 min, 22% B for 10 min. MS analysis was accomplished on Thermo Q-Exactive Plus Hybrid Quadrupole-Orbitrap Mass Spectrometer using electrospray ionization (ESI) in negative ionization mode. Samples were injected by a syringe pump at a flow rate of 5 μl‧min^-1^. The instrument parameters were as follows: spray voltage 3.5 kV, capillary temperature 320 ℃, sheath gas flow 30 mL‧min^-1^, aux gas flow 10 mL‧min^-1^, probe heated temperature 250 ℃ and m/z range 400-6000. Thermo Xcalibur Qual Browser was used for processing mass spectra and simulating isotopic distributions.

### AFM

The ASC protein was dissolved in NaOH solution at pH 7.5-8.0, and the final concentration was about 3.5 μM. The final concentration of Au_18_(L-NIBC)_14_ and Au_25_(L-NIBC)_18_ was 20 ppm. ASC alone or with either Au_18_(L-NIBC)_14_ or Au_25_(L-NIBC)_18_ are incubated in a water bath at 37 ℃. Sample was prepared by dripping 5.0 μL of solution of protein on freshly cleaved mica and allowed it to dry in the air. AFM experiments were carried on a FastScan Scanning Probe Microscopy (Bruker) with ScanAsyst in air mode at room temperature .

### DLS

DLS data were recorded on a Malvern Nano-ZS ZEN3600 zetasizer. 1.5 mL freshly prepared ASC solution (20 μg/mL) was added into a high-quality quartz glass cuvette for 52 h on-line continuous DLS measurement at 37 ℃. A working solution (1.5 mL) containing 20 μg/mL ASC and either Au_18_(L-NIBC)_14_ or Au_25_(L-NIBC)_18_ in ultrapure water with concentrations of 20 ppm was also added into a high-quality quartz glass cuvette for 52 h on-line continuous DLS measurement at 37 °C.

### RNA Sequencing Analysis

Cortical tissues were collected, and total RNA was extracted for sequencing. Quality control was performed using FastQC, followed by adapter trimming and low-quality read removal with Fastp. The cleaned reads were aligned to the reference genome using Hisat2, and gene expression levels were quantified with bowtie2. Differentially expressed genes (DEGs) were identified using DESeq2, applying a threshold of |log₂FC| > 1 and false discovery rate (FDR) < 0.05. A Venn diagram was generated to visualize overlapping and unique DEGs between the Naive *vs.* EAE and EAE *vs.* Au₁₈(L-NIBC)₁₄ comparisons. KEGG pathway enrichment analysis was performed, with key pathways such as the NOD-like receptor signaling pathway mapped via KEGG Mapper. The expression levels of selected genes were quantified using FPKM normalization, and statistical significance was assessed using student t-tests, with results visualized in a bar plot.

### MD simulations and analysis

All-atom MD simulations via GROMACS 2020.7 were conducted to probe the molecular interactions between ACS protein and gold nanoclusters using Charmm36 force field and TIP3P water model. The initial atom coordinate of ASC structure was extracted from the conformation of protein data bank (PDB) code 2KN6. The AuNCs nanostructures were generated and then combined through Material Studio software. The molar ratio of AuNCs:ASC is 1:1 in each MD system. Each MD system was placed in a periodic rectangular box and the periodic boundary conditions were applied to all of the xyz-directions. The MD were carried out via initial energy minimization, NVT equilibrium, NPT equilibrium and final production dynamics simulation. Specifically, the system’s heavy atoms were first constrained to minimize the energy of water molecules by 5000 steps with force constants of 1000 kJ·mol^-1^·nm^-2^; then, maintaining the constraints, 50000 step NVT ensemble simulation was carried out for the whole system. The temperature was 298 K, and the time step was 2 fs; Then 50000 step NPT ensemble simulation was carried out for the whole system, the temperature was 298 K, and the time step was 2 fs; Finally, the molecular dynamics simulation of the system was carried out in the NPT ensemble for 100 ns with a time step of 2 fs. During MD, the long-range interactions were processed using the PME method, while the van der Waals action was processed using the cut off method with a truncation value of 1.2 nm. The constant temperature was treated using the V-rescale method, and the air pressure was treated using the Berendsen method. An appropriate amount of Na and Cl ions were added to neutralize the ion systems. The equilibrium coordination and the underlying group interactions were analyzed via visual molecular dynamics (VMD).

### Cellular assay for inflammasome activation

Inflammasome activation was evaluated in PMA-differentiated THP-1 cells (100 ng/mL PMA for 48 h). The cells were stimulated with LPS (1 μg/mL) plus nigericin (20 μM) or poly (dA:dT) (2 μg/mL) or flagellin (1 μg/mL) or MDP (50 μg/mL) to generate inflammasomes NLRP3, AIM2, NLRC4 and NLRP1, respectively. Briefly, 1×10^6^ cells were plated in 12-well plates in 1 mL RPMI-1640 without supplements. Cells were primed for 3 h with 1 μg/ml of LPS, which were either mock-treated or primed with Au_18_(L-NIBC)_14_ or Au_25_(L-NIBC)_18_ during the last 30 min of LPS priming. To induce NLRP3 inflammasome, cells were further treated by nigericin for 30 min. To induce NLRC4 inflammasome, cells were stimulated for 3 h with flagellin. To induce AIM2 inflammasome, cells were transfected with poly (dA:dT) overnight using Lipofectamine 2000. To induce NLRP1 inflammasome, cells were stimulated for 2 h using MDP. Supernatants were harvested and clarified by centrifugation at 12000 rpm at RT, and IL- 1β and IL-18 releases were monitored by ELISA Kit following the manufacturer’s instructions^51,52^.

### MSU-induced peritonitis mouse model

C57BL/6 mice (9 weeks) were injected intraperitoneally with Au_18_(L-NIBC)_14_ or PBS (Naive group) 30 min before intraperitoneal injection of MSU (5 mg MSU crystals in 500 μl sterile PBS). 4 hours later, mice were sacrificed and peritoneal lavage with 5 ml PBS was performed, IL-1β secretion was detected by ELISA^53^.

### EAE induction and test of motor dysfunctions

All experimental procedures, including immunization, infusion, and decapitation of animals, were approved and performed according to the standards of the Ethics Committee for Animal Use of Wuhan University of Technology. All efforts were made to minimize the number of animals used and their suffering. EAE was induced on female C57BL/6 mice using Hooke Kit MOG35-55/CFA Emulsion PTX (Hooke laboratories, USA). A 100 μl emulsion containing myelin oligodendrocyte glycoprotein peptide fragment 35-55 (MOG35-55) in complete Freund’s adjuvant (CFA) was injected subcutaneously at each of two sites over the upper and lower back of mice, then injected with pertussis toxin intraperitoneally (i.p.), with the dose adjusted according to potency of each lot per manufacturer recommendations for each sex) on 0- and 1-day post-immunization. MCC950 (10 mg/kg), prednisone (10 mg/kg), Au_18_(L-NIBC)_14_ (20 mg/kg) or Au_25_(L-NIBC)_18_ (20 mg/kg) was injected i.p. once each day since day 0 (on day 0 and day 1, AuNCs were administrated after 1 hour of MOG35-55/CFA injection). Different dosages (1, 3, 10, 30 mg/kg) of Au_18_(L-NIBC)_14_ and oral administration of Au_18_(L-NIBC)_14_ (3 mg/kg) were also used to do the experiment to evaluate their influence on the pharmaceutical efficacy on the motor dysfunction of EAE mice. In above experiments, EAE scores for all mice were assessed every day to evaluate the clinical signs of mice: 0, no deficit; 0.5, partial tail limpness; 1, tail limpness; 2, tail limpness and hindlimb weakness; 2.5, paralysis of one hindlimb; 3, paralysis of both hindlimbs; 4, hindlimb and forelimb paralysis; 5, death^54,55^.

### Behavioral tests for early cognition and memory deficits of EAE mice

All the behavior tests for early cognition and memory deficits of EAE mice were performed between day 6 and day 13.

#### Open field test

In order to preclude the influence of motor behavioral difference on behavior tests for early cognition and memory deficits, the motor performances of mice were tested using open field test on day 9 after 3 days of adaption (day 6-8). On day 9, the mice were placed in the center of a 25 cm wide x 25 cm long x 20 cm high box. Prior to testing each animal, the entire open-field arena was cleaned using 75% alcohol. All open-field tests were 5 minutes in duration. The activity of mice in the box was digitally monitored and analyzed using Labmaze Animal Behavior Analysis Software V3.0 (Zhongshi dichuang technology, Beijing)^56^.

#### Barnes Maze

The mice were tested for spatial memory ability using a Barnes Maze (Zhongshi dichuang technology, Beijing) on day 13 after 3 days of adaption (day 6-8) and 4 days of training (day 9-12). On adaption days (day 6-8), the mice were placed in the center of the maze and subjected to aversive stimuli (bright light and noise). The mouse was given the opportunity to leave the maze through the escape hole. On training days (day 9-12), mice were subjected to 4 × 300 s trials per day (inter-trial interval time of 20 min). For probe trial on Day 13, mice were tested during a 300 s period of time. Animals were evaluated for their ability to remember the fixed position of an escape compartment. The time before reaching the target hole were recorded for the training and probe phases. For more consistency between animals, all training sessions took place between 7:00 and 12:00 a.m^57,58^.

#### Three-Chamber Sociability Test

Sociability and social memory were evaluated on a three-chamber sociability test on day 9 after 3 days of adaption (day 6-8). On adaption days (day 6-8), all mice were first habituated to an empty rectangle apparatus (Zhongshi dichuang technology, Beijing) for 5 min every day. The apparatus is divided into three compartments by two removable partitions with doors to connect adjacent two chambers. During the adaption day, the doors were kept open. On the testing day (day 9) the mice of Naive, EAE and Au_18_(L-NIBC)_14_ groups were firstly habituated to the central compartment for 10 min. Then, a strange mouse of the same strain, sex and age was placed in one of the two side compartments. The test mouse would socialize with this mouse for 10 min and it would become an “acquittance” to the test mouse. In the third stage, a new strange mouse of the same strain, sex and age (“stranger”) was placed in the compartment on the other side which is empty, and then the test mouse would socialize with the two mice at the same time for 10 min. The data were analyzed by comparing the time that the test mouse spent to get along with the “acquittance” mouse or the “stranger” mouse in the third stage^59,60^.

### H&E staining

On day 21 (the symptoms of EAE normally developed to the highest point^61^ on this day), the mice were anesthetized and perfused with 4% paraformaldehyde in PBS. The spinal cord was taken out and postfixed in 4% paraformaldehyde for 2 h, cryoprotected in 30% sucrose for 48 h, then embedded in a Tissue-Tek cryomold, and 10 μm sections were cut and collected on slides, stored at −80℃. Frozen sections of spinal cord were stained with hematoxylin and eosin (H&E) to assess immune cell infiltration. Slides were assessed in a blinded manner (by analysis of three fields)^62^.

### LFB staining

Paraffin embedded sections were incubated in 0.1% Luxol fast blue (LFB) overnight at 56°C, followed by washing using 95% ethanol and distilled water. Mount the sections on slides and cover them with a glass slide. Examine the stained sections under a microscope and capture images for further analysis. Blue areas indicate the intact myelin, whereas blank areas indicate demyelination. Demyelination was scored in a double-blinded fashion as followed: 0. no demyelination; 1. demyelination observed; 2. demyelination in several spots; 3. large area of demyelination^62^.

### Immunofluorescence staining

Frozen sections of spinal cord obtained by the above process were blocked with goat serum for 1 h at room temperature. Sections were incubated with Rabbit anti-MBP antibody (1:50) or Rabbit anti-ASC antibody (1:1000) overnight at 4℃. Then sections were washed and then incubated with secondary antibody conjugated to Alexa Fluor 568 (1:1,000) for 1 h at room temperature. Next, 10 μL of VectaShield Antifade Mounting Medium with DAPI was added to a glass slide. The stained sections were examined under a microscope, and both the DAPI-stained image and the MBP/ASC-stained image were captured for further analysis. In the images, MBP or ASC positive immunofluorescence staining areas were red, while those displaying DAPI positive immunofluorescence staining were blue. Subsequently, the DAPI-stained image and the MBP/ASC-stained image were merged. The proportion of MBP stained area in the white matter area of the spinal cord, as well as the ratio of cells with ASC specks, were calculated by ImageJ^63^.

### ELISA measurement

On day 21, spinal cords were harvested from mice and homogenized in RIPA Lysis Buffer using a motorized homogenizer. Subsequently, the samples were incubated on ice for 30 min and then centrifuged at 12,000 rpm for 15 min. The protein concentration was measured using the BCA Protein Assay. IL-1β, IL-18, TNF-α, Caspase-1, IL-17A and IFN-γ inflammatory cytokine levels were monitored by ELISA Kit following the manufacturer’s instructions.

### BBB permeability assay by Evans blue (EB) staining and FITC-dextran staining

The integrity of the BBB was examined using EB dye and 10kDa FITC-dextran as tracers. On day 30, mice were injected with 2% EB dye or 10kDa FITC-dextran (3 mg/mL) in PBS at a dosage of 2 mL/kg via the jugular vein. Two hours later, mice were transcardially perfused with PBS (200 mL) at a rate of 20 mL/min, then they were sacrificed immediately and their brains and spinal cords were collected. The wet weights of the organs were determined, and then the samples were homogenized in 3 mL of N, N-dimethylformamide. After incubation at 55°C for 24 h, the samples were centrifuged at 14,000 rpm for 20 min at 20°C. The supernatant was collected, and the absorbance was measured at 620 nm (EB) or 520 nm (10kDa FITC-dextran) using a Jenaway 6300 spectrophotometer^64,65^.

### Western blot for occludin

On day 30, the mice were sacrificed and microvessels from mice spinal cords were isolated and homogenized in cold sucrose solution (0.32 M Sucrose, 5 mM HEPES, pH 7.4). The samples were centrifuged at 1000×g for 10 min at 4 ℃, and the pellets were suspended in cold sucrose solution. This suspension was centrifuged at 350×g for 10 min at 4 ℃ and the supernatant was discarded. Then the pellets were lysed in RIPA buffer. After centrifugation at 3000×g for 15 min at 4°C, the supernatant was collected. The protein concentration of the supernatant was determined using the BCA Protein Assay. After adding loading buffer to the supernatant and mixing well, the protein was fully denatured in a metal bath at 100 ℃ for 5 min. Thirty micrograms of denatured protein were separated by SDS-PAGE gels and transferred to nitrocellulose membranes. The membranes were incubated with Mouse anti-Occludin antibody (1:1000) or GAPDH (1:1000) overnight at 4°C. Then, the membranes were washed and incubated with the corresponding HRP-conjugated goat anti-mouse secondary antibody (1:1000) for 1h at room temperature. The bands were detected by ECL chemiluminescence kits and the gray value of the bands was analyzed using ImageJ software^66^.

### DSS-Induced Colitis Model

Female C57BL/6 mice (8–10 weeks old, 18–21 g) were randomly assigned to four groups (n=10 per group), consisting of one Naive control group and three experimental groups. The experimental groups were provided with 3% DSS in their drinking water from Day0 to Day6 to induce colitis, while the Naive group received regular drinking water. Test compounds (2 mg/kg and 6 mg/kg) were administrated intraperitoneally from Day0 to Day7. The vehicle was administrated intraperitoneally with saline during the same period of time. Body weight and DAI scores were recorded daily to monitor disease progression. On Day 8, the mice were euthanized, and colon lengths were measured to evaluate the extent of inflammation and tissue damage.

### TNBS-Induced Colitis Model

Female C57BL/6 mice (8–10 weeks old, 18–21 g) were randomly divided into four groups (n=10 per group). The Naive group received a rectal administration of 100 µL of 50% ethanol, while the three experimental groups were administered 100 µL of 1.5% TNBS in 50% ethanol to induce colitis. Test compounds (3 mg/kg and 6 mg/kg) were administered intraperitoneally from Day-1 to Day4. The vehicle was administrated intraperitoneally with saline during the same period of time. Body weight and DAI scores were recorded daily. On Day 5, the mice were euthanized, and colon lengths were measured to assess the degree of inflammation.

### IMQ-induced psoriasis model

Female C57BL/6J mice (6–7 weeks old) were randomly allocated into four groups (n=10 per group), comprising one Naive control group and three experimental groups. A 2 cm × 3 cm area on the dorsal skin of the mice was shaved. Two hours later, the experimental groups received a topical application of approximately 50 mg of 5% imiquimod (IMQ) cream on the back and 5 mg on both sides of the right ear, while the Naive group was treated with petroleum jelly. These treatments were applied once daily for nine consecutive days. From Day 0 to Day 8, test compounds and vehicle were administered via intraperitoneal injection. PASI scores and right ear thickness were measured daily to evaluate disease progression and therapeutic efficacy.

### Statistical Analysis

All data are presented as mean ± standard error of the mean (SEM)unless otherwise stated. Statistical analyses were performed using GraphPad Prism 9 software. Comparisons between two groups were made using unpaired two-tailed Student’s t-test. For comparisons among multiple groups, one-way analysis of variance (ANOVA) followed by Dunnett’s Multiple Comparisons Test was used. All experiments were performed at least three times independently.

---

**Extended Data Fig. 1.**
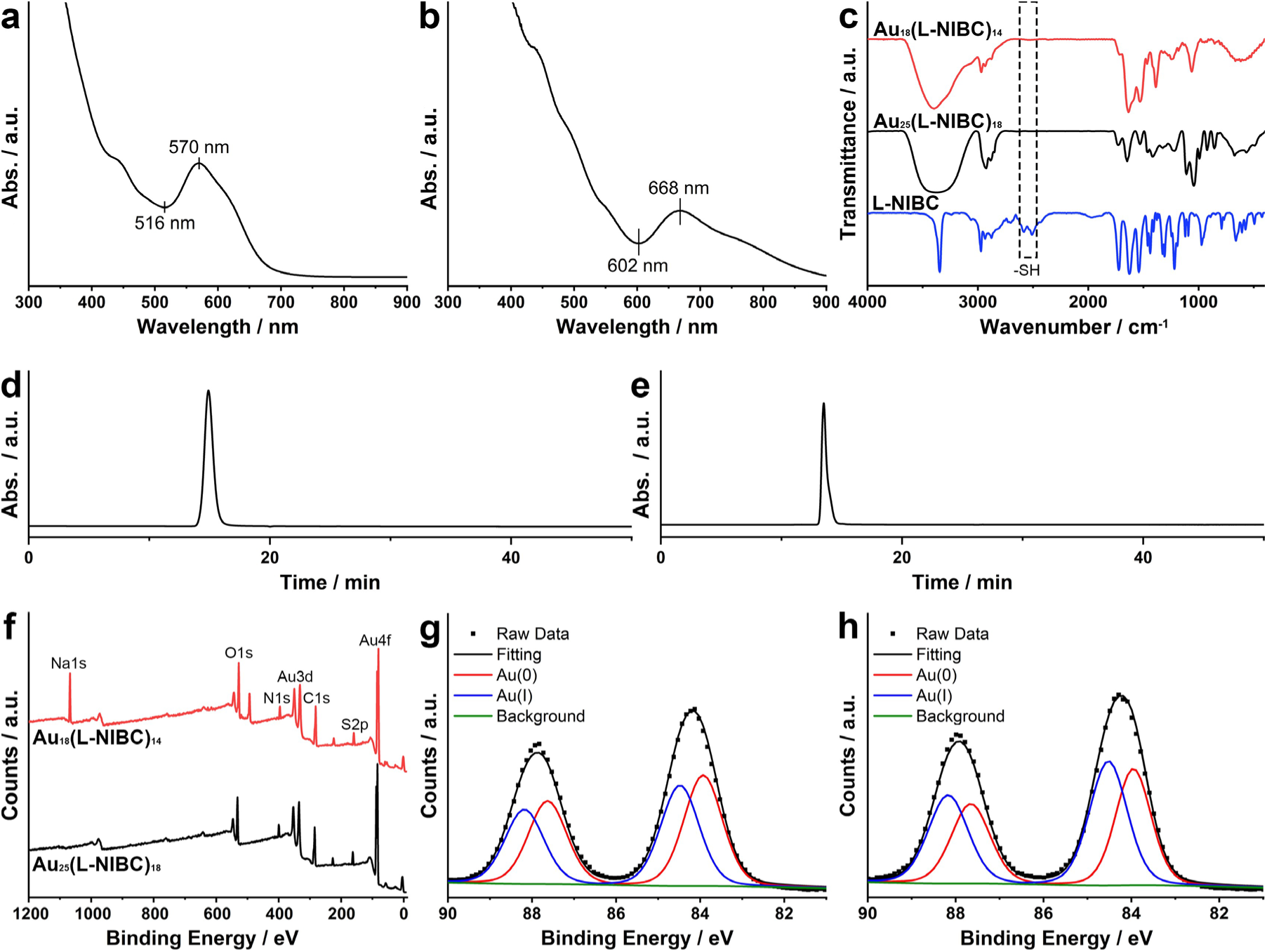
Other characterization data for two AuNCs sample. **a**, **b**, UV- Vis absorption spectra. **c**, Infrared spectra. **d**, **e**, SEC chromatograms. **f**, Full-scan XPS spectra. **g**, **h**, High-resolution Au4f XPS spectra. **a**, **d**, **g**, Au_18_(L-NIBC)_14_. **b**, **e**, **h**, Au_25_(L-NIBC)_18_.

**Extended Data Fig. 2.**
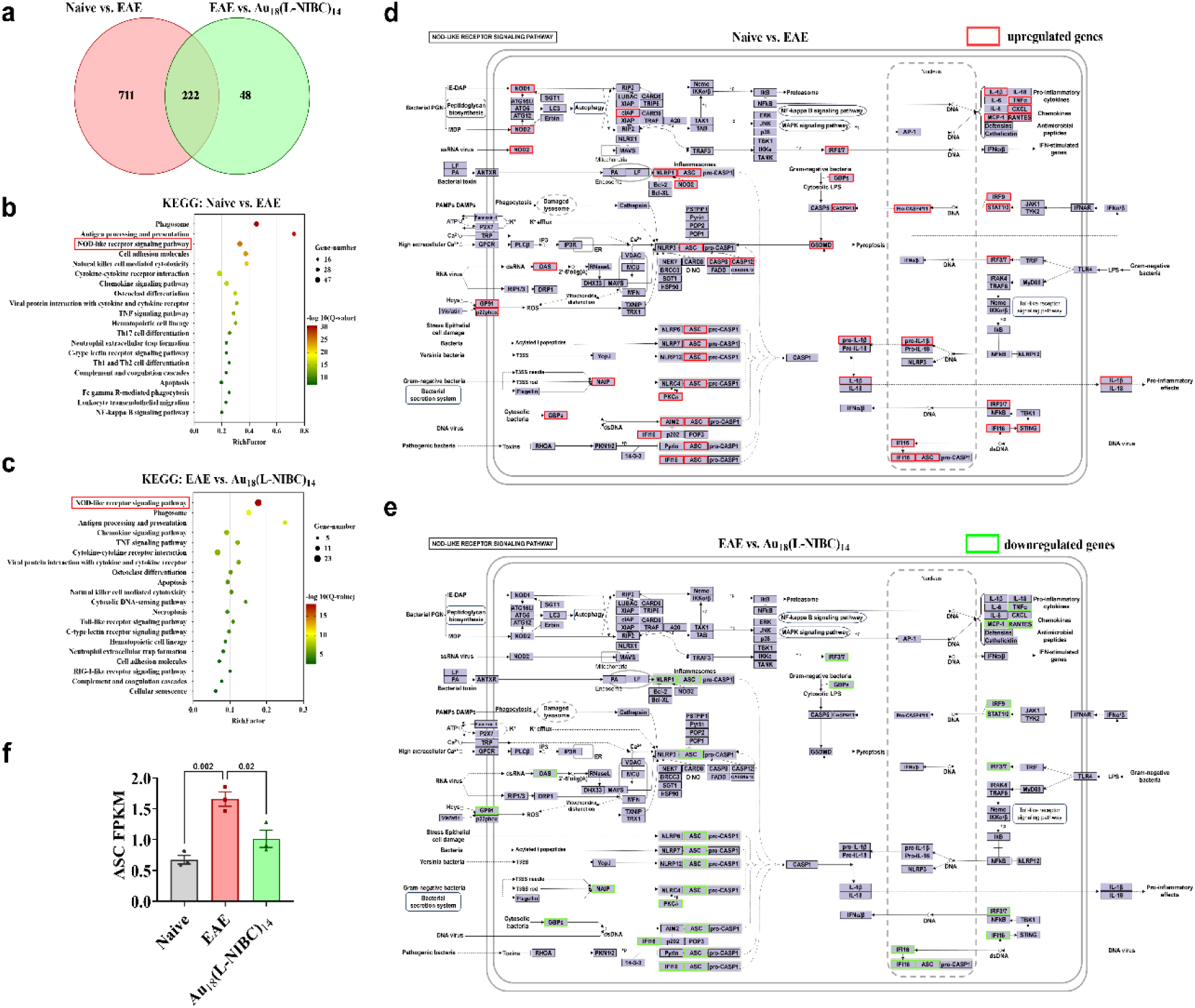
RNA-Seq analysis of cortical tissues from Naive mice, EAE mice, and Au₁₈(L-NIBC)₁₄-treated EAE mice. **a**, Venn diagram showing differentially expressed genes among the three groups. **b**, **c**, KEGG pathway enrichment analysis for Naive *vs.* EAE (**b**) and EAE *vs.* Au₁₈(L-NIBC)₁₄ (**c**). **d**, **e**, Pathway diagrams highlighting upregulated (red) and downregulated (green) genes in the NOD-like receptor signaling pathway for Naive *vs.* EAE (**d**) and EAE *vs.* Au₁₈(L-NIBC)₁₄ (**e**). **f**, Comparison of ASC fragments per kilobase of transcript per million mapped reads (FPKM) across the three groups, with statistical significance indicated.

**Extended Data Fig. 3.**
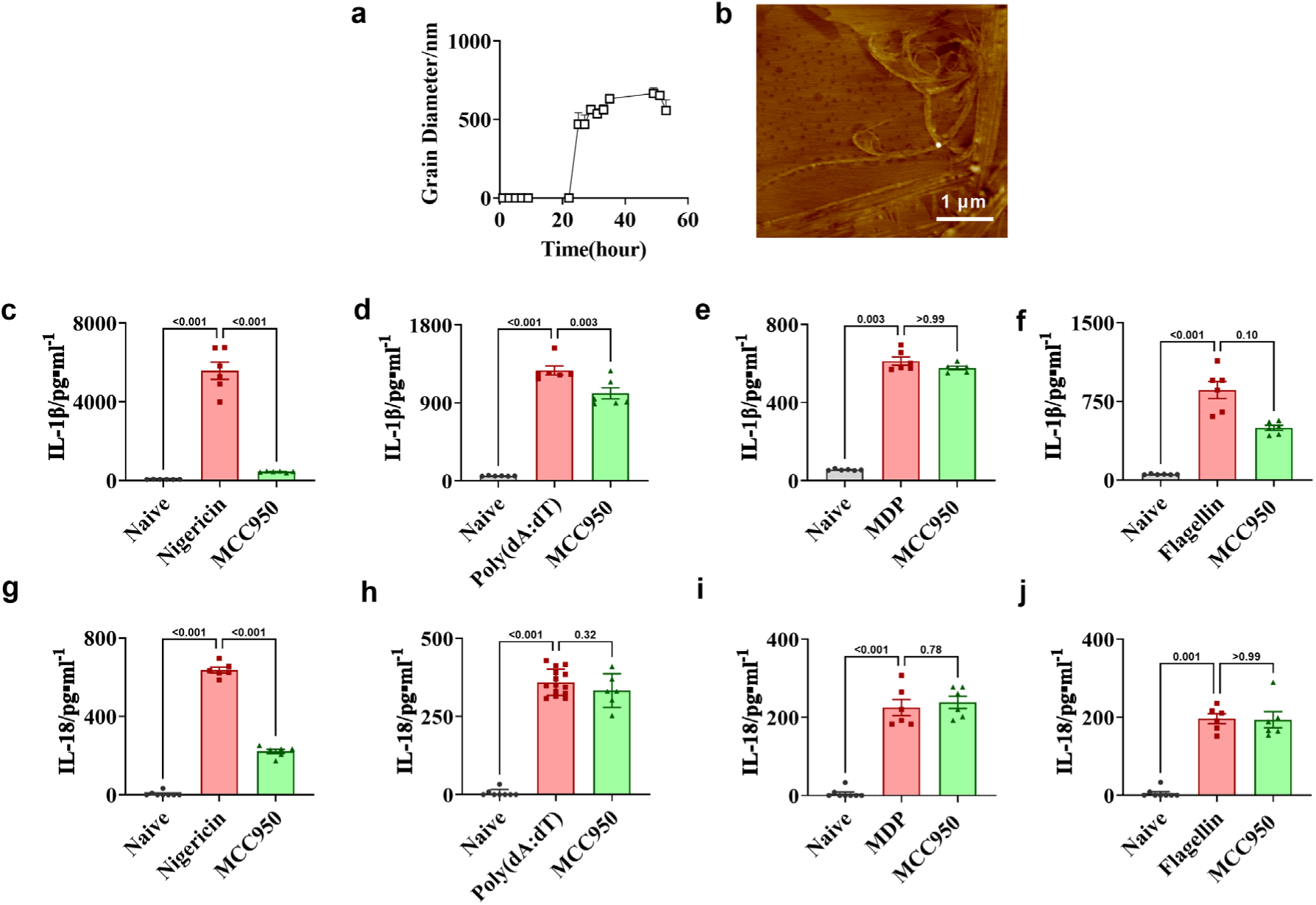
MCC950 is a specific inhibitor for NLRP3 which cannot inhibit ASC oligomerization directly. DLS experiment (**a**) shows that coincubation with MCC950 did not postpone the ASC fibrillation time. AFM image after coincubation for 48 hours (**b**) indicates the formation of thick ASC fibers which is similar to ASC incubation alone. Cell experiments using PMA differentiated THP-1 cells stimulated by LPS plus nigericin (**c**, **g**), dA:dT (**d**, **h**), MDP (**e**, **i**), and flagellin (**f**, **j**) show that MCC950 can significantly inhibit the activation of NLRP3 inflammasome, but is ineffective to other inflammasomes.

**Extended Data Fig. 4.**
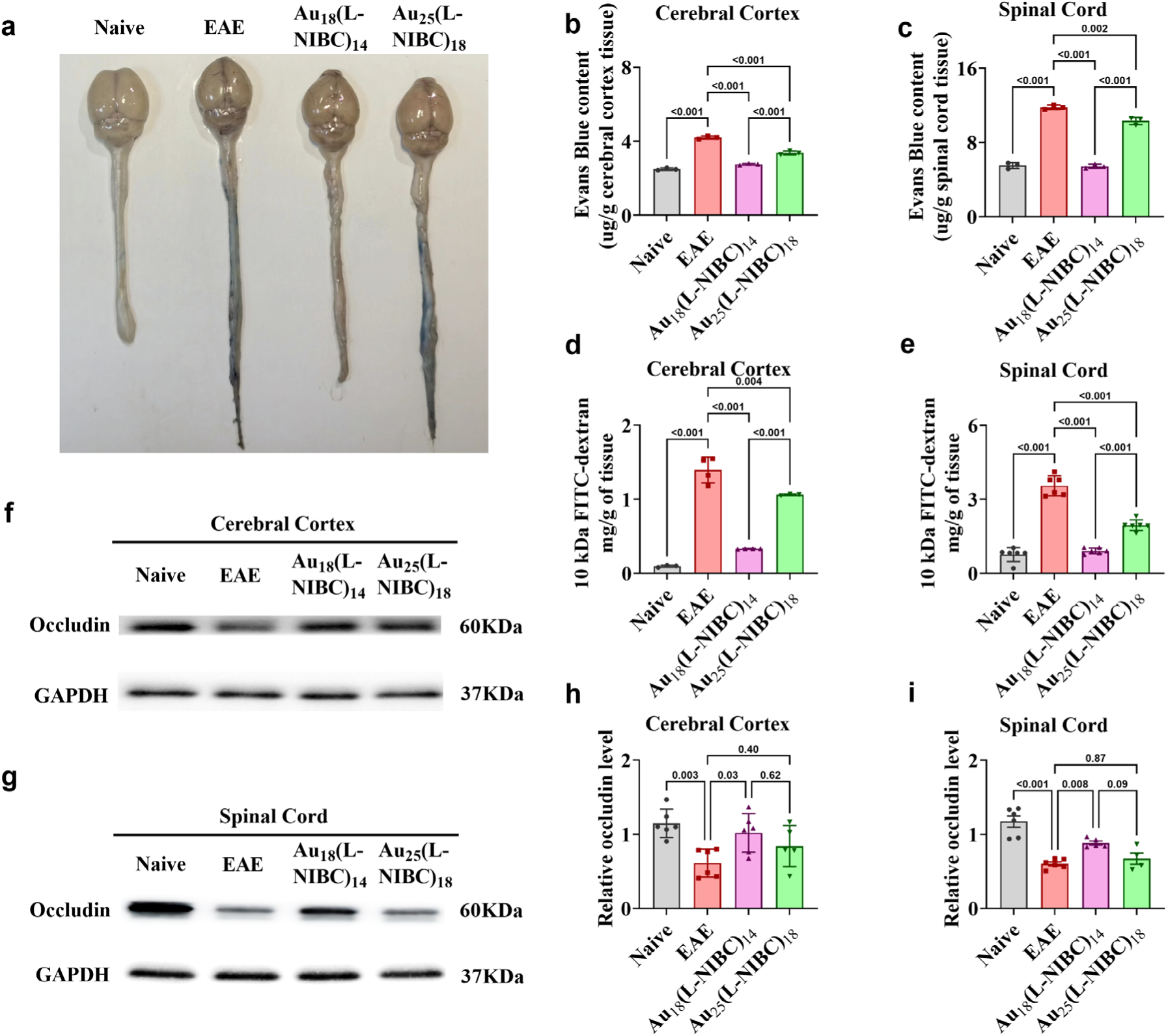
Au_18_(L-NIBC)_14_ shows excellent BBB protection properties. **a**, representative images of EB dye extravasation in the brains and spinal cords of EAE mice with or without treatments of Au_18_(L-NIBC)_14_ or Au_25_(L-NIBC)_18_. **b** and **c**, quantitative results of EB dye extravasation experiment in cerebral cortices (**b**) and spinal cords (**c**) of mice. **d** and **e**, quantitative results of FITC-dextran staining experiment in cerebral cortices (**d**) and spinal cords (**e**) of mice. **f** and **g**, representative WB patterns of occludin in cerebral cortices (**f**) and spinal cords (**g**) of mice. **h** and **i**, expressions of occludin protein normalized to GAPDH in cerebral cortices (**h**) and spinal cords (**i**) of mice.

**Extended Data Fig. 5.**
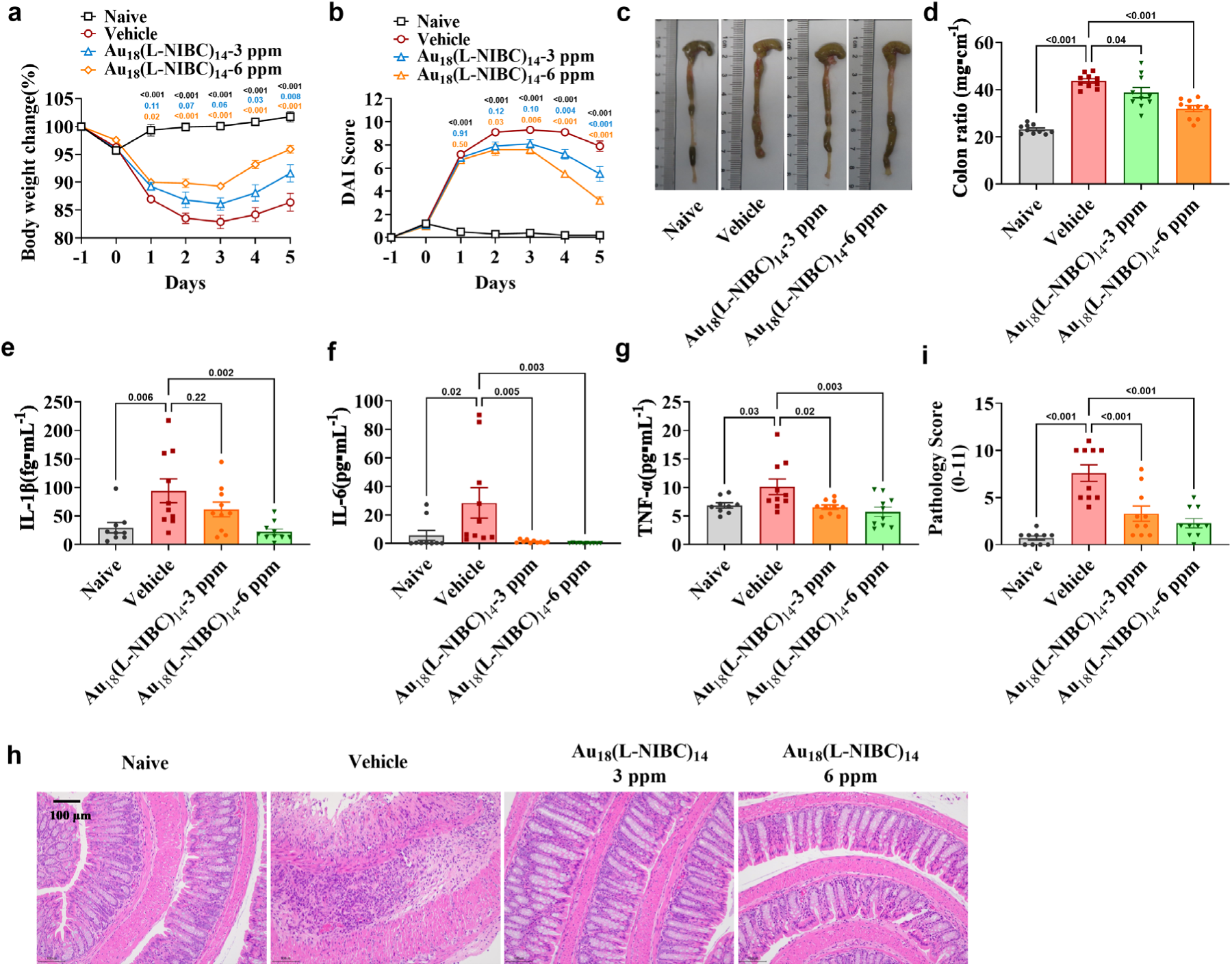
Therapeutic effects of Au_18_(L-NIBC)_14_ on TNBS induced colitis mouse model. **a**, body weight change. **b**, DAI score. **c**, representative images of colon showing the colon lengths. **d**, colon ratio. **e-g**, concentrations of IL-1β (**e**), IL-6 (**f**), and TNF-α (**g**) in serum. **h**, representative images of H&E staining of colon slices. **i**, pathological scores of colon tissues obtained from H&E staining.

**Extended Data Fig. 6.**
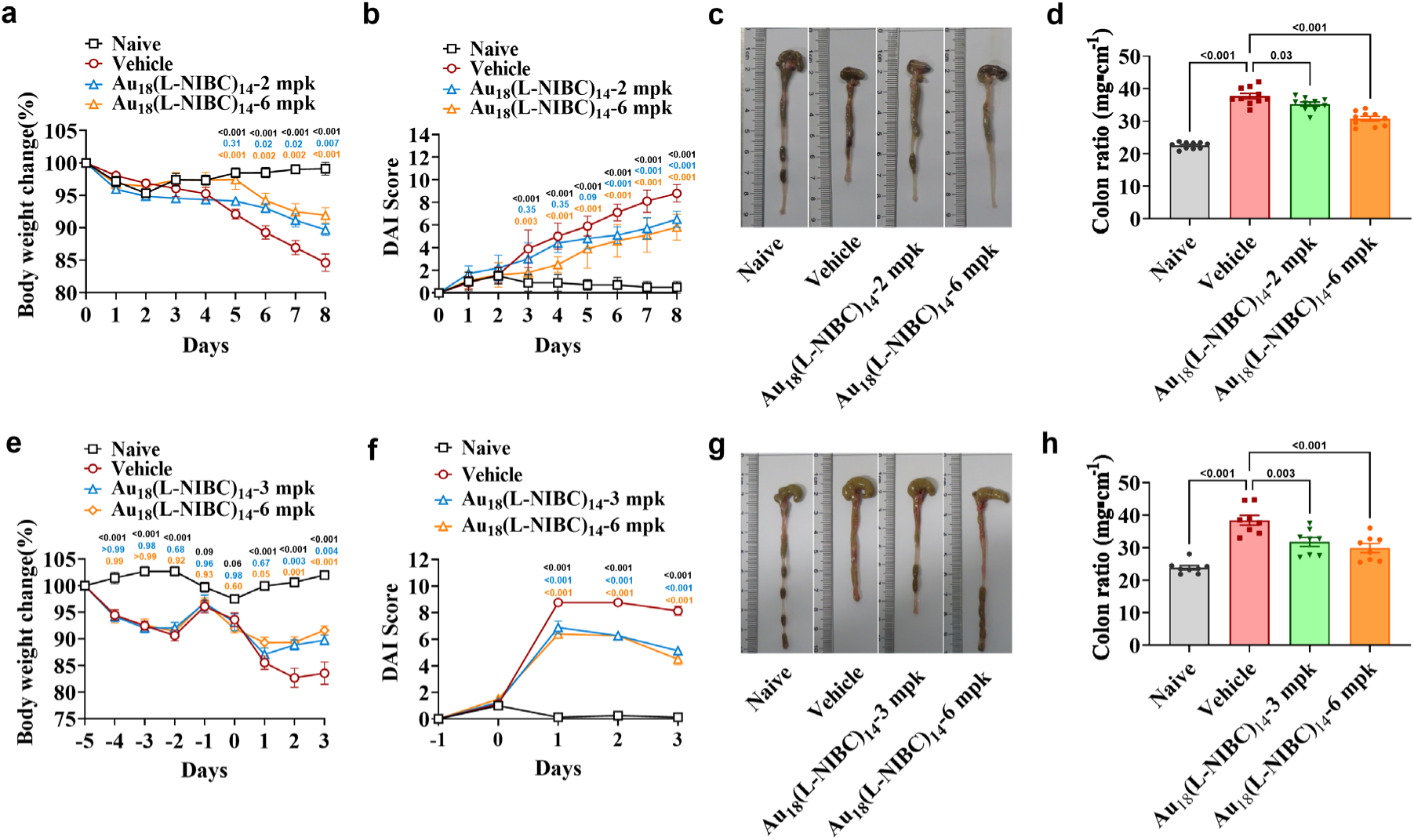
Therapeutic effects of Au_18_(L-NIBC)_14_ on DSS-induced and OXZ-induced colitis mouse models. **a-d**, DSS-induced colitis model. **e-h**, OXZ-induced colitis model. **a** and **e**, body weight change. **b** and **f**, DAI score. **c** and **g**, representative images of colon showing the colon lengths. **d** and **h**, colon ratio.

**Extended Data Fig. 7.**
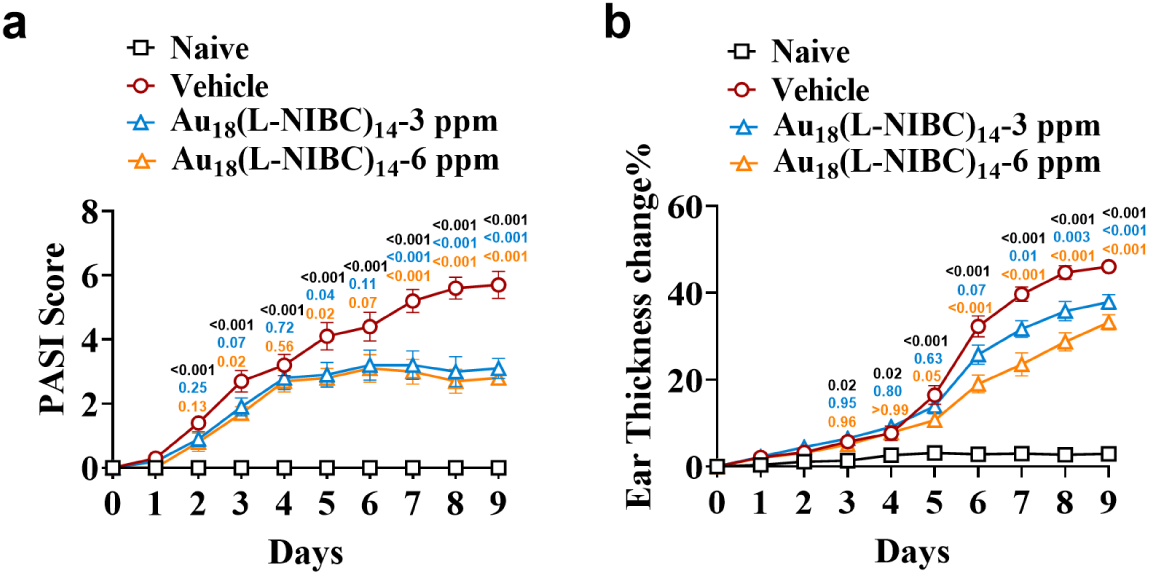
Therapeutic effects of Au_18_(L-NIBC)_14_ in IMQ-induced psoriasis mouse model. **a**, PASI score. **b**, ear thickness change.

**Extended Data Fig. 8.**
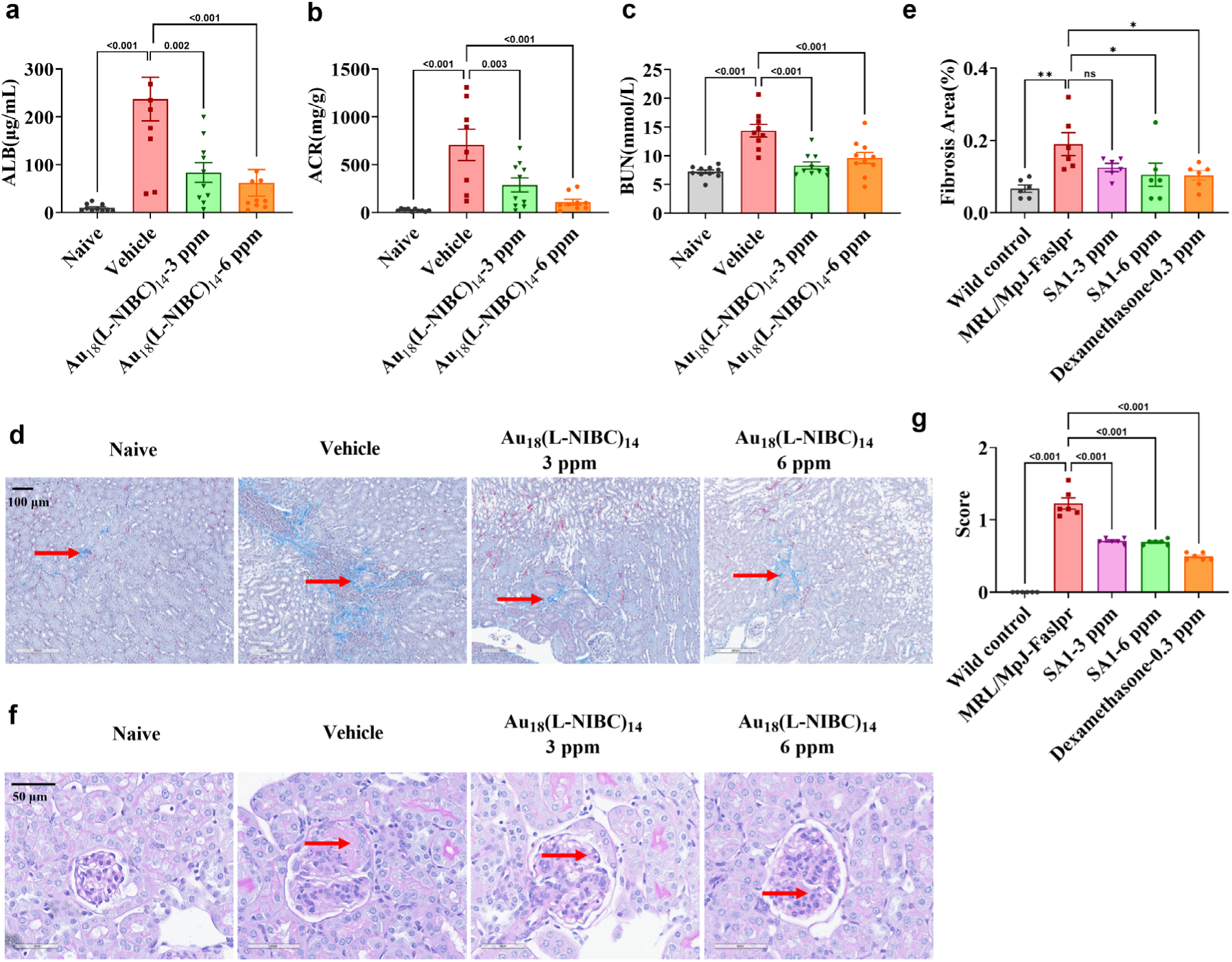
Therapeutic effects of Au_18_(L-NIBC)_14_ in female MRL/MpJ-Faslpr mouse model. **a**, urine albumin (ALB) concentration. **b**, the ratio of urine albumin to creatinine (ACR). **c**, blood urea nitrogen (BUN) concentration. **d**, representative images for M&T staining showing the renal fibrosis. **e**, renal fibrosis areas. **f**, representative images for periodic acid-Schiff (PAS) staining showing the glomerulosclerosis. **g**, glomerulosclerosis index (PASI score).

## Notes

### Competing Interest Statement

The authors have declared no competing interest.

